# Site-specific methionine oxidation alters structure and phase separation of TDP-43 C-terminal domain

**DOI:** 10.64898/2025.12.15.694486

**Authors:** Busra Ozguney, Ryan Z. Puterbaugh, Renjith Viswanathan, Jayakrishna Shenoy, Priyesh Mohanty, Jeetain Mittal, Nicolas L. Fawzi

## Abstract

TAR DNA binding protein 43 (TDP-43), a key protein linked to ALS pathology, undergoes phase separation and forms functional assemblies via condensation within cells. The conserved region (CR) within its C-terminal domain (CTD) mediates self-assembly through helix-helix interactions, while the flanking intrinsically disordered regions (IDRs) contribute to phase separation through transient interactions involving aromatic and hydrophobic residues. The CTD contains ten methionine residues distributed equally between these regions, making it particularly susceptible to oxidative modifications. While methionine oxidation is known to impair phase separation, neither the precise mechanism nor the specific contribution of methionines in the CR compared to the IDRs has been determined. Here, we combine NMR spectroscopy and all-atom molecular dynamics (MD) simulations to reveal if and how methionine oxidation in each region differentially affects CTD structure and phase separation. We demonstrate that all methionine residues are vulnerable to oxidation, leading to distinct regional effects: oxidation of CR methionines disrupts helical structure and directly impairs intermolecular helical association, while oxidation of IDR methionines disrupts long-range contacts. Hence, oxidation of methionines in both regions contributes to impaired phase separation, albeit through different mechanisms. These findings establish methionines as critical redox-sensitive modulators of TDP-43 phase behavior and provide molecular insights into how oxidative stress may contribute to TDP-43 dysregulation in neurodegenerative diseases.

## INTRODUCTION

Eukaryotic cells organize their biochemical processes through membraneless organelles (MLOs), including nucleoli, stress granules (SGs), and various ribonucleoprotein (RNP) granules^1^. These biomolecular condensates form via phase separation, where proteins, nucleic acids, and other molecules spontaneously de-mix into concentrated liquid droplets through weak, multivalent interactions^2–5^. Beyond simple compartmentalization, phase separation orchestrates transcriptional regulation, stress response, and synaptic signaling^6^. However, this dynamic organization via transient interactions and without a bounding membrane can also make these condensates highly responsive to cellular conditions, mutations, and post-translational modifications (PTMs) ^7–12^. This sensitivity can lead to dysregulation of biomolecular condensates in diseases, termed *condensatopathies*, which are increasingly implicated in neurodegeneration, cancer, viral infections, and cardiomyopathy^13^. While cells normally regulate condensate formation through multiple mechanisms including temperature, pH, chaperone interactions, and cellular modifications^13^, disruption of these regulatory mechanisms can trigger pathological transitions^10^. This phenomenon is especially evident in neurodegenerative diseases where RNA-binding proteins (RBPs) containing prion-like domains (PrLDs) mediate both normal physiological phase separation and pathological aggregation^14^. Among these RBPs, TAR DNA binding protein 43 (TDP-43) has emerged as a central factor in diseases like amyotrophic lateral sclerosis (ALS) and frontotemporal dementia (FTD), where its aberrant aggregation is a hallmark pathological feature^15^.

TDP-43 is a 414-residue multi-domain protein that shuttles between nucleus and cytoplasm, playing crucial roles in mRNA metabolism, localization and transport^16–18^. Its architecture includes a folded N-terminal domain (NTD), two evolutionary conserved RNA-recognition motifs (RRM1 and RRM2), and a C-terminal glycine rich low-complexity domain (CTD) ^19^. TDP-43 CTD holds significant disease relevance as it harbors the majority of the disease-associated mutations and forms pathological aggregates in patient neurons^20–23^. TDP-43 CTD comprises intrinsically disordered regions (IDR1, aa: N267-I318; glutamine/asparagine (Q/N) rich region, aa: S342-Q365; and IDR2, aa: A366-M414) flanking a hydrophobic segment that adopts a transient α-helical structure (conserved region, CR aa: N319-A341) that mediates helical multimerization^24–28^. Under physiological conditions, TDP-43 CTD readily undergoes phase separation in the presence of salt or RNA^24^. The integrity of this helical multimerization is vital for this function, as demonstrated by disease-associated mutations that disrupt helical stability or interactions and impair phase separation^24,29–31^. While phase separation plays important regulatory roles in TDP-43 function, clustering of TDP-43 may also serve as a precursor to pathological aggregation. Under cellular stress conditions, TDP-43 is recruited to cytoplasmic stress granules, creating environments of high local protein concentration^31,32^. This prolonged residence in a phase-separated state may promote aberrant transitions from liquid droplets to more solid-like aggregates and eventually to pathological fibrils^33–35^.

Oxidative stress is strongly implicated in aging^36^ and neurodegenerative diseases including ALS and Alzheimer’s disease^37^, introducing another layer of complexity to TDP-43 regulation. The brain is particularly vulnerable to oxidative damage due to its high metabolic activity and relatively low levels of antioxidant enzymes^38–42^. This vulnerability increases with age as antioxidant defense systems become less efficient, leading to the accumulation of oxidative modifications to proteins, lipids, and nucleic acids^43^. At the protein level, oxidation can occur through various mechanisms, including modifications of specific amino acids such as methionine (Met), cysteine (Cys), tryptophan (Trp), and tyrosine (Tyr) ^44–46^. Among these, modification of Met residues has been associated with modulation of protein self-assembly and amyloid fibril formation^47–53^.

Compared to most IDRs, TDP-43 CTD is distinctively enriched in Met residues, distributed across the CR and the flanking IDRs (**Figure 1A, Figure S1A).** Notably, Met residues comprise nearly 25% of the CR (5 of 21 residues). Methionine enrichment in TDP-43 CTD is evolutionarily conserved among vertebrates, with TDP-43 homologs maintaining similar Met distribution across both CR and CR-flanking disordered regions (**Figure S1A).** Furthermore, interactions involving these residues are crucial for phase separation as demonstrated by mutation studies where methionine substitution with alanine significantly reduce condensate formation^54^. Upon oxidation of the thioether group to sulfoxide^55^ (**Figure 1B),** the resulting increase in the side chain polarity ^44,45^ may disrupt protein structure and Met-mediated interactions^56^, although precise mechanism remains unclear.

**Figure 1.**
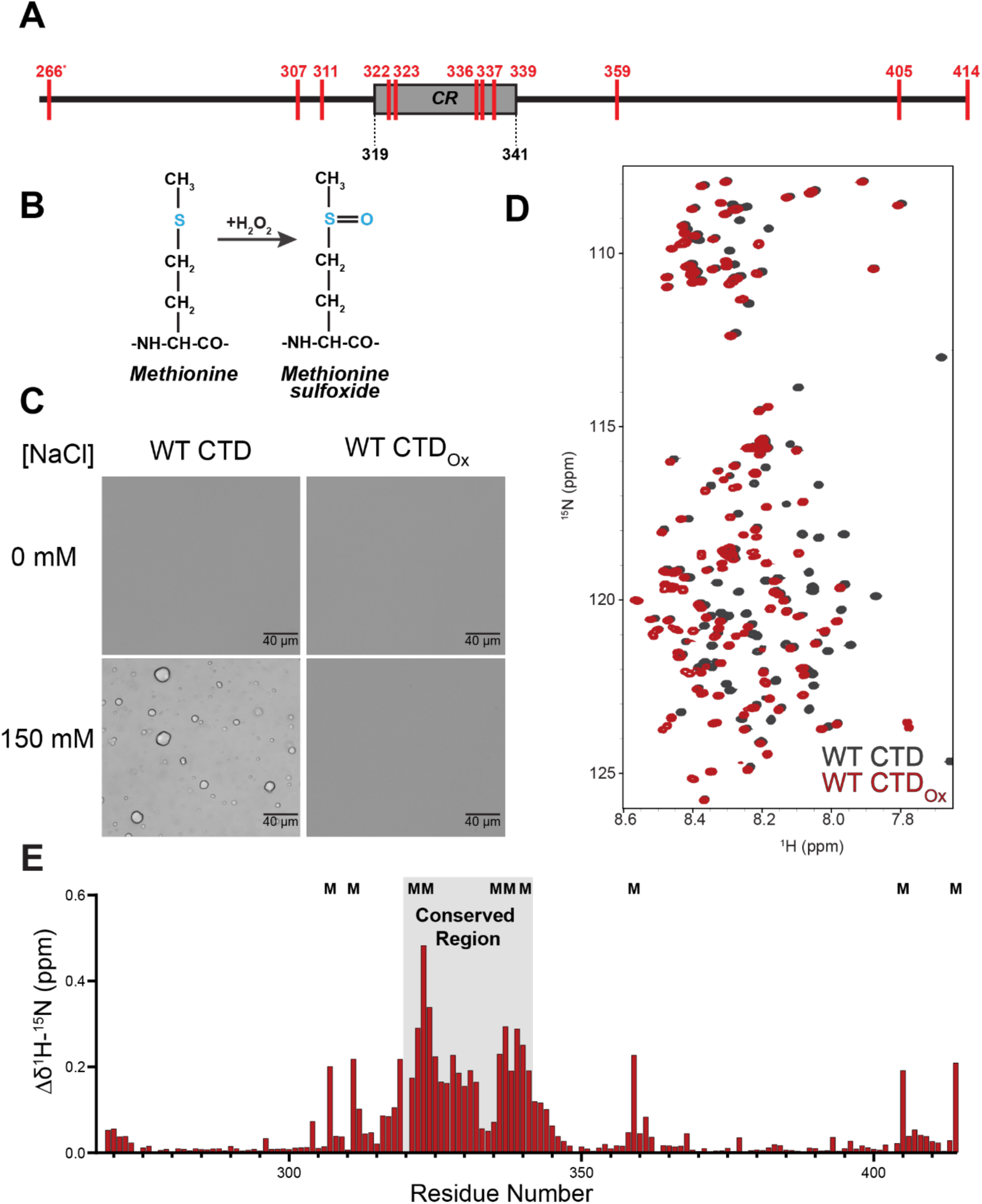
Oxidation of methionine residues changes the structure of TDP-43 CTD and disrupts phase separation. **(A)** Schematic illustration of the TDP-43 C-terminal domain (CTD), showing the positions of methionine residues (highlighted in red) and the conserved region (CR) which is critical for protein-protein interactions and phase separation. **(B)** Diagram depicting the conversion of a methionine residue (left) to methionine sulfoxide (right) via oxidation of the methionine’s thioether group with hydrogen peroxide. **(C)** DIC micrographs of both the unoxidized and oxidized WT TDP-43 CTD in the presence and absence of 150 mM NaCl (80 μM protein, 20 mM MES pH 6.1) demonstrating that increasing ionic strength results in the formation of liquid-like protein droplets by the unoxidized TDP-43 CTD while the oxidized form remains in the dispersed phase. **(D)** 1H-15N HSQC fingerprint spectra of unoxidized (grey) and oxidized (red) CTD (90 μM protein, 20 mM MES pH 6.1, 360 mM Urea) displaying dramatic shifting of residue backbone peaks particularly in the CR region of the protein. **(E)** Combined ^1^H-^15^N CSPs of WT CTD following oxidation demonstrate that conserved region residues undergo the largest CSPs following oxidation. Methionines in the disordered region also undergo large CSPs but this effect is isolated mainly to the methionine themselves with flanking residues experiencing weaker shifts.

Indeed, recent studies investigating the impact of oxidative stress on TDP-43 CTD have yielded conflicting results. Lin et al.^57^ proposed that a cross-β structure in the CR formed under physiological conditions provided greater protection against oxidation for two of its methionines (M322/323) compared to other positions, for both *in vitro* samples (hydrogels and droplets) and within cells. Conversely, Carrasco et al.^58^ found no evidence of β-structure and demonstrated that oxidation disrupts CR helicity, leading to complete disruption of phase separation upon oxidation. Thus, the specific impact of Met oxidation on TDP-43 CTD structural integrity and phase separation remains incompletely understood. Furthermore, whether the position of Met residues within the CR versus the IDRs differentially influences their role as redox modulators of TDP-43 function is unclear. To address these questions and resolve existing discrepancies, we investigated how methionine location affects phase separation and self-assembly under oxidizing conditions using complementary experimental and computational approaches.

Through a combination of nuclear magnetic resonance (NMR) experiments and all-atom molecular dynamics (AAMD) simulations, we examine the effects of oxidation on protein structure and interactions in a position-dependent manner. Using biochemical experiments, we separate and probe the contributions of IDR and CR methionine oxidation on the driving forces for phase separation. Furthermore, we complement our experimental findings with atomistic simulations, revealing that TDP-43 phase separation depends on both structural integrity of the CR helix and a network of Met-mediated contacts, including interactions with both polar (Q, N, S, R), aliphatic (A, L, M), and aromatic (F, W) residues, that are disrupted upon oxidation. These insights provide a molecular framework for understanding how oxidative stress may contribute to the TDP-43 dysfunction at the molecular level.

## RESULTS

### Methionine oxidation of CTD disrupts phase separation and reduces CR helicity

We first investigated the impact of oxidation on CTD phase separation and structure. Treatment of TDP-43 CTD with 1% H_2_O_2_ has been previously demonstrated to oxidize the naturally occurring thioether group of Met to sulfoxide without progressing to a sulfone ^54^. Indeed, we observed using mass spectrometry that treatment of the CTD with 1% H_2_O_2_ for 30 mins resulted in the predominant species within our samples displaying a ∼176 Da increase in mass corresponding to the addition of 11 oxygen atoms confirming that all Met residues (comprising 10 from the native CTD sequence and 1 from an N-terminal cloning artifact) had been oxidized to sulfoxides (**Figure S1B**). DIC microscopy revealed that the while the unmodified CTD readily undergoes phase separation in the presence of NaCl, oxidation of Met residues results in the CTD losing the ability to undergo phase separation under the same conditions (**Figure 1C**). Importantly, we performed these studies by pre-oxidation and then full removal of the oxidizing agent to control for any effect of the hydrogen peroxide on the solution conditions, demonstrating the loss of phase separation is solely due to the oxidation of Met residues. Interestingly, NMR fingerprint spectra (^1^H-^15^N heteronuclear single quantum coherence, HSQC) of TDP-43 CTD before and after oxidation (CTD_ox_) (**Figure 1D**) displayed numerous chemical shift perturbations (CSPs) demonstrating that oxidation changed the chemical environment of residues beyond just the Met residues alone. Mapping CSPs as a function or residue position demonstrated that the largest CSPs corresponded to residues within the Met-rich CR (**Figure 1E**). IDR Met residues also displayed large CSPs, however, unlike the CR, these CSPs were mostly confined to the Met residues themselves with adjacent residues displaying much weaker CSPs. Collecting ^1^H-^13^C HSQCs of the CTD before and after oxidation also displayed large CSPs (**Figure S2A**), with the largest corresponding to the Met side chain carbons. The CTD_ox_ spectrum displayed no evidence of unmodified Met residues, demonstrating complete oxidation at these reaction conditions regardless of sequence position.

We then tested the effect of TDP-43 oxidation on the multimerization of TDP-43 and the mechanistic reason for this effect. Previous NMR and simulation studies have demonstrated that the CR contains a short α-helical structure that mediates the formation of helical assemblies and CTD multimers^25,31^. The self-assembly of the CTD is enhanced with increasing protein concentration, which results in CR residues displaying CSPs when comparing CTD NMR fingerprint spectra at low and high concentrations^24^. When we collected spectra of the CTD at a low (20 μM) and high (90 μM) concentration, we observed strong CSPs localized to the CR, as previously demonstrated. When we repeated this experiment with CTD_ox_ we observed no large CSPs (**Figure 2A-B**). These data indicate that oxidation of the CR Met residues appears to largely disrupt the formation of CTD self-assemblies. The ^1^H-^15^N HSQC fingerprint spectrum of CTD_ox_ displayed a narrower peak dispersion than the CTD (**Figure 1D**), an indication of reduced structural order, consistent with prior reports that oxidation of the CR Met residues disrupts the helicity of the region^58^. To provide further evidence of this disruption, we determined the secondary chemical shift (ΔδC_α_-δC_β_) for CR residues before and after oxidation. We observed that before oxidation, resonances of Ala residues within the CR display a strong positive value, indicative of a α-helix (**Figure 2C-D**). But following oxidation of neighboring Met residues, these resonances shift in position closer to those of a random coil, confirming that oxidation reduces the α-helicity of this region. Overall, our findings demonstrate that all Met residues in CTD are susceptible to oxidation, leading to significant disruption of helical structure and self-assembly.

**Figure 2.**
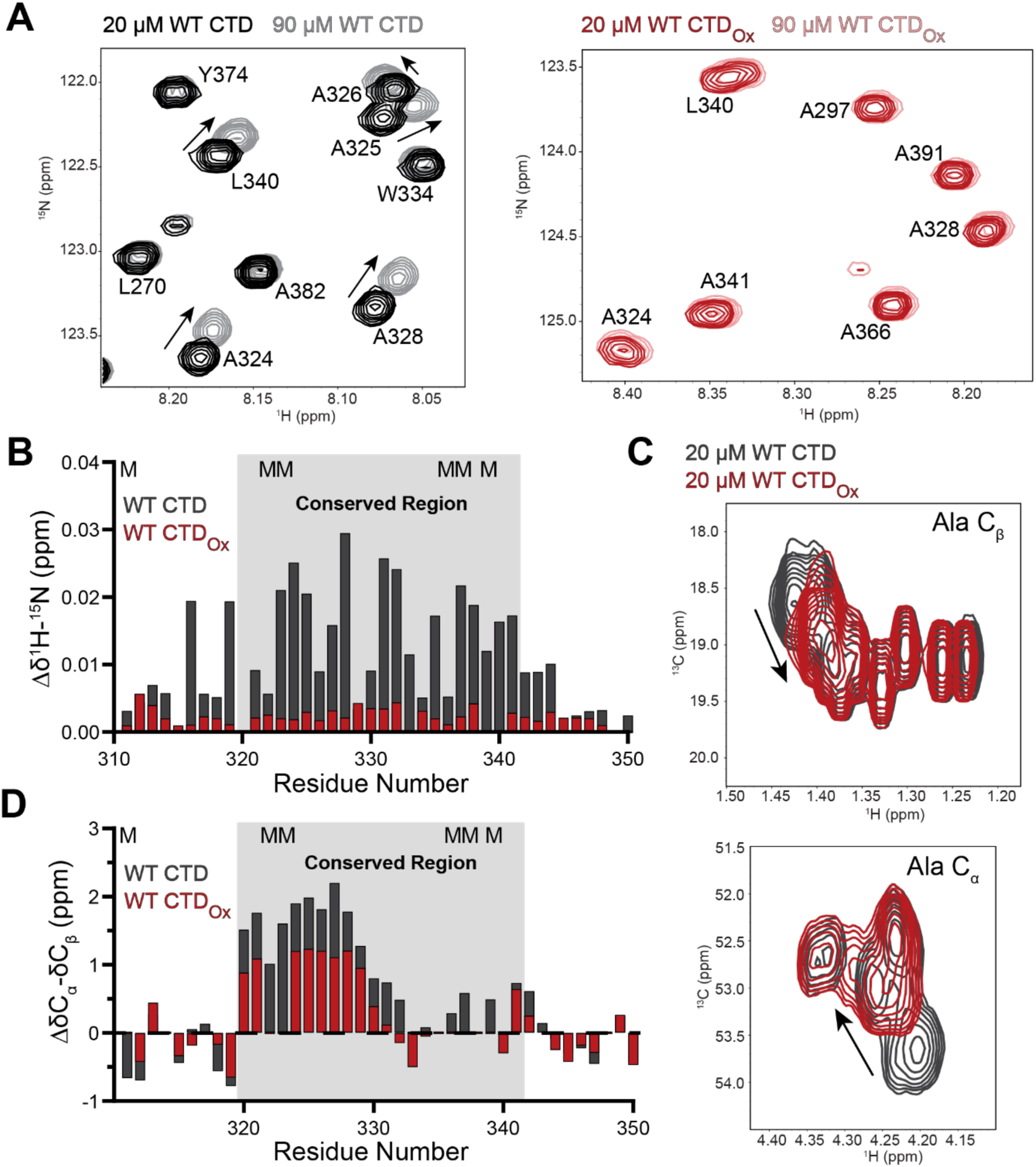
Oxidation of methionine residues reduces cooperative helical assembly of the CTD by reducing the helicity of the conserved region. **(A)** Overlay of ^1^H-^15^N HSQC spectra of 20 μM (black) and 90 μM (grey) unoxidized WT CTD and 20 μM (red) and 90 μM (pink) oxidized WT CTD demonstrating that chemical shift perturbations (CSPs) of CR residues due to increased helical assembly at higher concentrations is disrupted by oxidation of methionine residues. **(B)** Combined ^1^H-^15^N CSPs of unoxidized (grey) and oxidized (red) CTD demonstrate that the magnitude CSPs across the CR are significantly reduced by the oxidation of methionine residues. This demonstrates that the oxidation of methionine residues prevents the formation of cooperative helical assembly by the CR. **(C)** Overlay of ^1^H-^13^C HSQC spectra of 20 μM unoxidized WT CTD (grey) and 20 μM oxidized WT CTD (red) demonstrating that the carbon chemical shift of alanine residues within the CR undergo significant CSPs following oxidation. **(D)** Secondary carbon chemical shifts within the CR of the unoxidized WT CTD (grey) and oxidized WT CTD (red) display a decrease in positive CR secondary shifts following oxidation. This demonstrates that the helicity of the region is reduced by the oxidation of methionine CR residues explaining the loss of cooperative helical assembly by this region.

### Methionine oxidation promotes expansion of the CTD monomer and disrupts interactions involving polar, aliphatic, and aromatic residues

Our experimental findings demonstrated that oxidation of all Met residues in the CTD significantly disrupts helical structure and impairs phase separation. However, the atomic-level details of how Met oxidation induces these changes remain unclear. We and others have previously established that the extent of chain compaction for IDPs at dilute concentrations positively correlates with their ability to undergo LLPS at higher concentrations^54,59–62^. Further, for disordered low-complexity domains containing repetitive sequence regions, intramolecular (residue-level) interactions observed in single-chain AAMD simulations are strongly correlated with intermolecular interactions implicated in their phase separation^59,63,64^. Based on these established correlations, we analyzed the conformational ensembles of unoxidized CTD and its methionine oxidized variant, CTD_ox_, generated from multi-microsecond AAMD simulation trajectories to obtain mechanistic insights into how oxidation impacts the CTD secondary structure, conformations, and intramolecular interactions disrupting its phase separation (see **Figure S3**, **Material and Methods**).

Consistent with NMR experiments, the CTD_ox_ ensemble displayed a significant disruption of α-helicity in the CR region compared to the unoxidized ensemble (**Figure 3A-B, S4 and S5**). Further, comparison of the radius of gyration (R_g_) probability distributions revealed that the CTD_ox_ ensemble adopted more expanded conformations compared to unoxidized CTD (**Figure 3C),** consistent with the inability to undergo phase separation. Analysis of root mean square intrachain distances (R_ij_) as a function of sequence separation (|i-j|) showed that chain expansion is more pronounced for residues with sequence separation exceeding 40 residues, indicating disruption of long-range interactions (**Figure 3C inset)**. Importantly, disruption of long-range interactions in repetitive IDRs like those in TDP-43 CTD also correlates with disruption of homotypic phase separation^54,63^. To understand the basis of conformational expansion upon oxidation, we further analyzed the intramolecular interaction network based on the averaged pairwise contact maps (over three replicas) for both CTD and CTD_ox_ ensembles, capturing any heavy atom interaction pairs within 4.5 Å between residues separated by at least two positions (|i-j|>3). We found that upon oxidation, the interaction landscape becomes more diffuse with significantly lower contact formed compared to the unoxidized ensemble (**Figure S6A**). Furthermore, we quantified interactions within and between CTD regions (IDR1, CR, Q/N, IDR2) by summing the contacts between all possible pairs throughout the simulations and analyzed the resulting distributions. The analysis revealed that most inter-region interactions are disrupted upon Met oxidation (**Figure S6B**).

**Figure 3.**
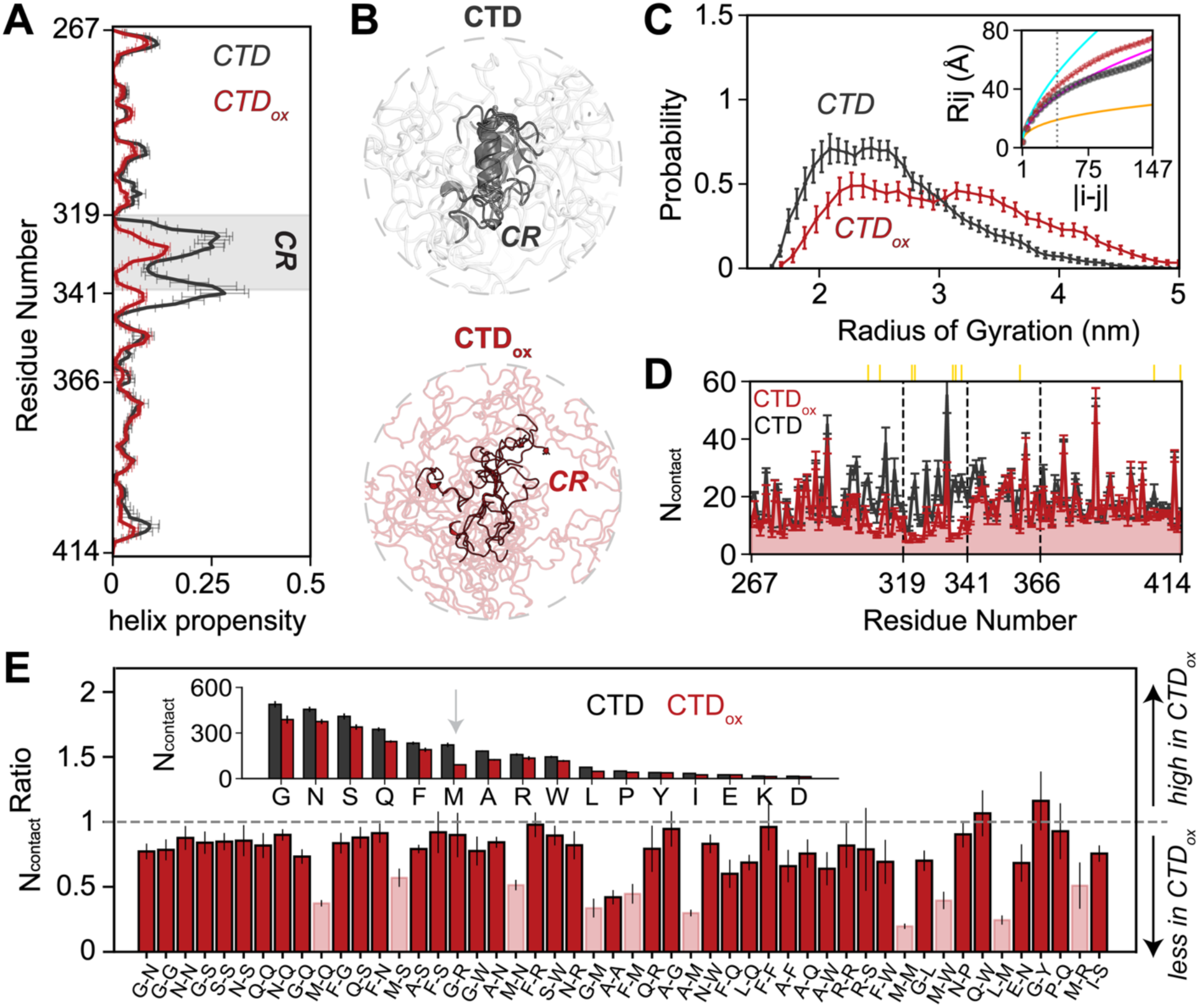
Impact of methionine oxidation on the conformational ensemble of the TDP-43 C-terminal domain (CTD). **(A)** Per-residue helical propensity analysis revealing a disruption of secondary structure in the CR region upon methionine oxidation. **(B)** Representative structural ensembles of the CTD and CTD_ox_ proteins. Oxidation of the methionine residues within and around the CR disrupting its secondary structure, essentially causing the helix to melt into a more flexible coil. **(C)** Radius of gyration (R_g_) distributions comparing the overall compactness of wild-type (CTD, gray) and methionine-oxidized (CTD_ox_, red) TDP-43 CTD. The shift toward larger *R*_g_ in the CTD_ox_ protein indicates that oxidation causes an expansion of the CTD. Inset: Root mean square intrachain distance (R_ij_) as a function of sequence separation (|i-j|). Reference scaling laws for ideal chain (magenta, ν=1/2), self-avoiding random walk (cyan, ν=3/5), and collapsed globule (yellow, ν=1/3) are shown, with b=5.5 Å. **(D)** One-dimensional projection of the average residue-residue contact maps, quantifying the total contacts per residue. **(E)** Quantification of contacts between specific residue types. Ratio is calculated by dividing number of contacts for each residue pair in CTD_ox_ with CTD. Number of contacts formed by each residue type is shown in the inset. Methionine oxidation reduces contacts for most residue pairs. Error bars when shown represent the standard error of the mean across three independent replicas for CTD and CTD_ox_. Together, these data demonstrate that oxidation of methionine residues in the TDP-43 CTD causes an expansion of the disordered protein, disrupts its conserved helical structure, and alters its intramolecular interaction patterns.

To assess residue-level contributions to the weaker intramolecular interactions upon oxidation, we summed all pairwise contacts for each residue position. This analysis showed that the reduction in pairwise contacts for CTD_ox_ was most pronounced at Met-containing positions (**Figure 3D**), particularly in regions with high Met density (C-terminus of IDR1 and CR regions). Further comparison of the top 50 residue-type contact pairs (**Figure S7 and 3E**) revealed that contacts involving Met residues showed the largest decrease in CTD_ox_. Polar (Q, N, S, R), aliphatic (A, L, M), and aromatic (F, W) residues exhibited significant decreases (nearly two-fold or more) in their interactions with Met, as evident from the contact ratios computed relative to the unoxidized ensemble (**Figure 3E**). Overall, our results indicate that the CTD interaction network undergoes a significant reorganization upon oxidation, with loss of intramolecular contacts involving polar, aliphatic, and aromatic residues and reduced CR helicity. These changes lead to conformational expansion, which is linked to CTD’s inability to undergo phase separation, as observed in experiments.

### Oxidation of methionine impairs CTD phase separation regardless of position but disrupts CR helicity only upon CR oxidation

Our previous work^54^ shed light on the importance of Met residues in promoting CTD phase separation through their contribution to the IDR-mediated interactions increasing the CTD multivalency. We found that substituting Met residues in both the CR and the flanking IDRs with alanine, which has a higher helical propensity but lower hydrophobicity, impaired phase separation and increased the saturation concentration (C_sat_), defined as the minimum concentration at which droplet formation is observed, compared to wild-type (WT). In contrast, replacing IDR Met residues with leucine (L), which has similar hydrophobicity to Met, maintained phase separation behavior comparable to WT. Intriguingly, substitution with aromatic tyrosine (Tyr) enhanced phase separation and decreased C_sat_^54^. These substitution studies demonstrated that altering the identity of residues at Met positions impacts phase separation, with different mechanisms depending on the domain location of the modification. Since Met oxidation similarly reduces hydrophobicity by converting the thioether group to a polar sulfoxide group, we next tested whether oxidation in different regions (CR or IDRs) would have distinct effects on phase separation.

To gain a deeper understanding of the role of Met oxidation in different regions, we designed two CTD variants: (1) 5M→A^IDR^, a variant where all five Met residues in the IDRs are substituted with Ala and therefore only contains Met residues in the structured CR region, and (2) 5M→A^CR^+ 5A→M^IDR^, a variant where all five Met residues in the CR are replaced with Ala, and five Ala residues in the IDRs are replaced with Met and therefore only contains Met residues in the unstructured IDR regions (**Figure S8A**). The latter variant was designed because removing the CR Met residues (i.e. 5M→A^CR^) prevents phase separation at accessible concentrations^54^, and this variant swapping Met positions (5M→A^CR^+ 5A→M^IDR^) is reasonable as it does preserve the overall amino acid composition of TDP-43 CTD. DIC microscopy revealed that both variants undergo phase separation in the presence of NaCl although we did observe a difference in droplet size, with the 5M→A^IDR^ variant forming much larger droplets on average than the 5M→A^CR^+ 5A→M^IDR^ variant, likely due to the lower C_sat_ for 5M→A^IDR^, which we previously explored ^54^ (**Figure 4A**). Despite this difference, following oxidation, both variants displayed a complete loss of phase separation. These findings demonstrate that oxidation disrupts Met interactions not only in the structured CR but also in the unstructured CR-flanking regions, and that disrupting interactions in both regions contributes to a loss of phase separation.

**Figure 4.**
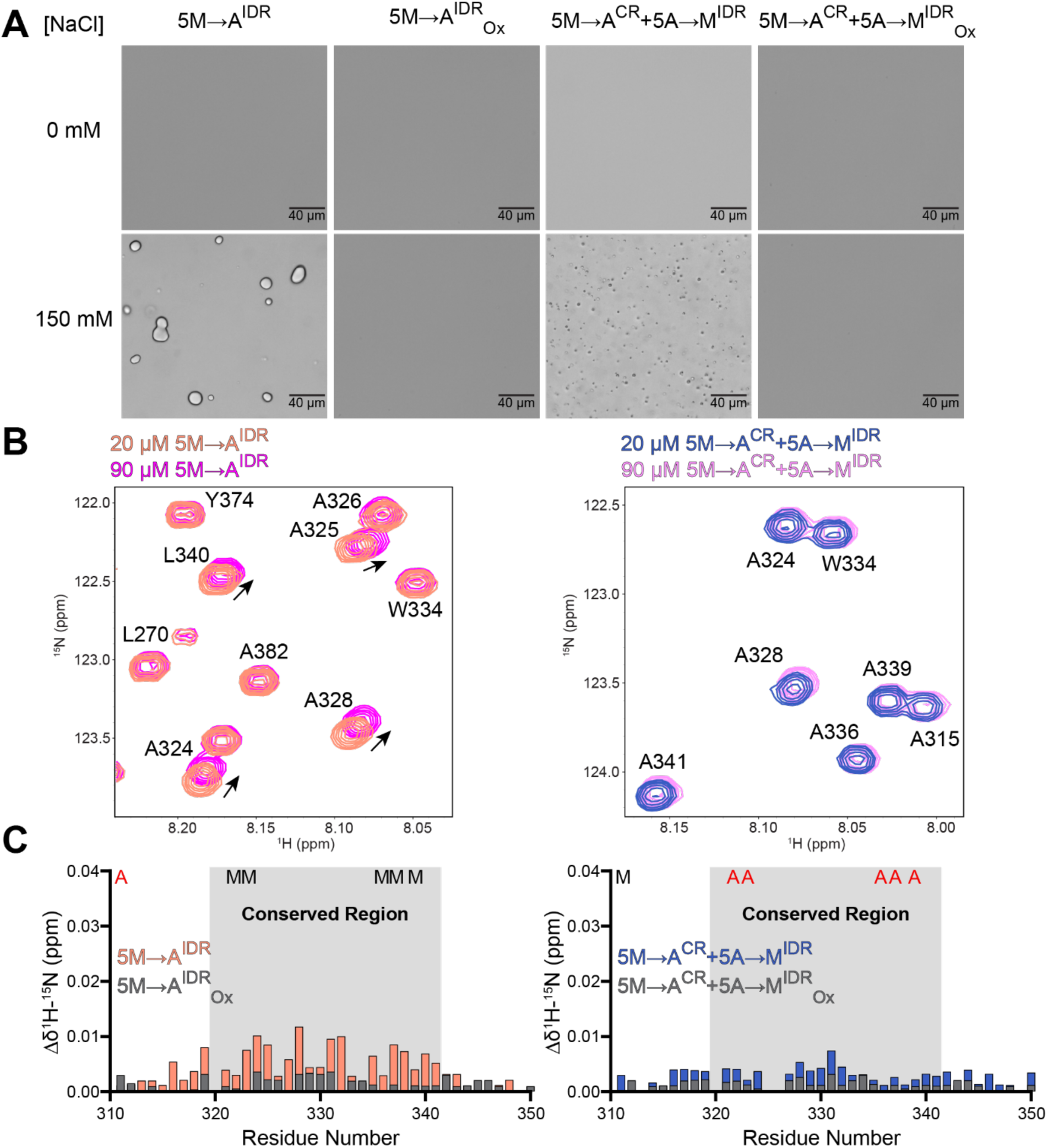
Oxidation of methionine residues within the conserved helical region and the disordered flanking regions both contribute to disrupting phase separation of TDP-43 CTD. **(A)** DIC micrographs of both the unoxidized and oxidized 5M→A^IDR^ and 5M→A^CR^+5A→M^IDR^ CTD in the presence and absence of 150 mM NaCl (80 μM protein, 20 mM MES pH 6.1). Increasing ionic strength results in the formation of large liquid-like droplets by 5M→A^IDR^ similar to the WT, while 5M→A^CR^+5A→M^IDR^ formed smaller droplets. Oxidation of both constructs resulted in the loss of phase separation. This demonstrates that oxidation of methionine residues in both the helical region and disordered regions contribute to the disruption of PS. **(B)** Overlay of ^1^H-^15^N HSQC spectra of 20 μM (salmon) and 90 μM (magenta) unoxidized 5M→A^IDR^ CTD and 20 μM (blue) and 90 μM (pink) unoxidized 5M→A^CR^+5A→M^IDR^ CTD displaying that 5M→A^IDR^ CR residues display significantly stronger CSPs than 5M→A^CR^+5A→M^IDR^ due to the loss of CR methionine. This demonstrates that IDR methionines in 5M→A^CR^+5A→M^IDR^ mediate contacts important for phase separation. **(C)** Combined ^1^H-^15^N CSPs of both unoxidized (salmon) and oxidized (grey) 5M→A^IDR^ CTD and unoxidized (blue) and oxidized (grey) 5M→A^CR^+5A→M^IDR^ CTD displaying a greater decrease in CR residue CSPs following oxidation of 5M→A^IDR^ than 5M→A^CR^+5A→M^IDR^. This demonstrates that the CR region of 5M→A^CR^+5A→M^IDR^ is relatively unaffected by oxidation further demonstrating that loss of PS is due to disruption in disordered region interactions.

To provide additional evidence that Met residues interactions in disordered regions are disrupted by oxidation, we acquired NMR spectra of the 5M→A^IDR^ and 5M→A^CR^+ 5A→M^IDR^ variant, both unoxidized and oxidized, at a high (90 μM) and low (20 μM) concentration (**Figure 4B, Figure S8B**). We observed that the unoxidized 5M→A^IDR^ variant displayed large CSPs within the CR region (**Figure 4B-C**), albeit weaker than the WT, demonstrating this variant formed helical assemblies qualitatively similar to the WT. Again, similar to the WT, the oxidized form of the 5M→A^IDR^ variant displayed no strong CSPs demonstrating the oxidation once again disrupted assembly mediated by the helical CR region. Conversely the 5M→A^CR^+ 5A→M^IDR^ variant displayed very small CSPs even before oxidation (**Figure 4B-C**), demonstrating that this variant does not strongly form helical assemblies and therefore interactions are mediated primarily within the disordered regions. Additionally, we observed the peak positions of CR residues did not drastically change when comparing the oxidized and unoxidized 5M→A^CR^+ 5A→M^IDR^ variant (**Figure 4B, Figure S8B**), suggesting that helicity is retained following oxidation in the absence of Met residues, demonstrating that disruption of phase separation for this variant that lacks Met in the CR is not due to loss of helicity in the CR region. We also utilized a third variant 5M→A^CR^, a variant where all five Met residues in the CRs are substituted with Ala, to investigate how oxidation might affect picosecond-nanosecond motions near IDR Met residues by measuring ^15^N transverse relaxation (*R_2_*). We found that while the majority of the residues retained similar *R_2_* values before and after oxidation, there was a difference in the values of M307 and M311 and their surrounding residues (**Figure S8C**). This demonstrates that oxidation does have a minor effect on molecular motions of IDR Met residues and their immediate surroundings, providing further evidence that oxidation disrupts Met interactions in the IDRs. Overall, these data demonstrate that oxidation does not just disrupt phase separation by disrupting the helicity and helix-helix interactions of the CTD’s CR, but also by disrupting Met interactions within the IDR, both of which contribute the loss of phase separation.

### Effect of position-specific oxidation on secondary structure, conformations, and CTD self-association

Selective oxidation of Met residues in individual CTD regions is not achievable experimentally, as all accessible Met residues undergo oxidation. This precludes direct investigation of region-specific oxidation effects on phase separation without substantially altering the amino acid sequence (e.g., 5M→A^CR^ + 5A→M^IDR^). To overcome this limitation and complement our experimental findings, we next investigated the consequences of Met oxidation in specific CTD regions on the monomer ensemble by analyzing the AAMD-derived ensembles of CTD variants with partial oxidation (CR_ox_ and IDR_ox_) (**Figure S9**). Our analysis revealed distinct structural consequences of region-specific oxidation (**Figure 5, S10-11, S12A**). CR oxidation caused a dramatic reduction in α-helicity to levels seen in CTD_ox_, whereas IDR oxidation preserved the helical structure. While both variants showed expansion of the CTD upon Met oxidation, CR_ox_ was notably more expanded than IDR_ox_, to levels similar to CTD_ox_ (**Figure S12B-C**).

**Figure 5.**
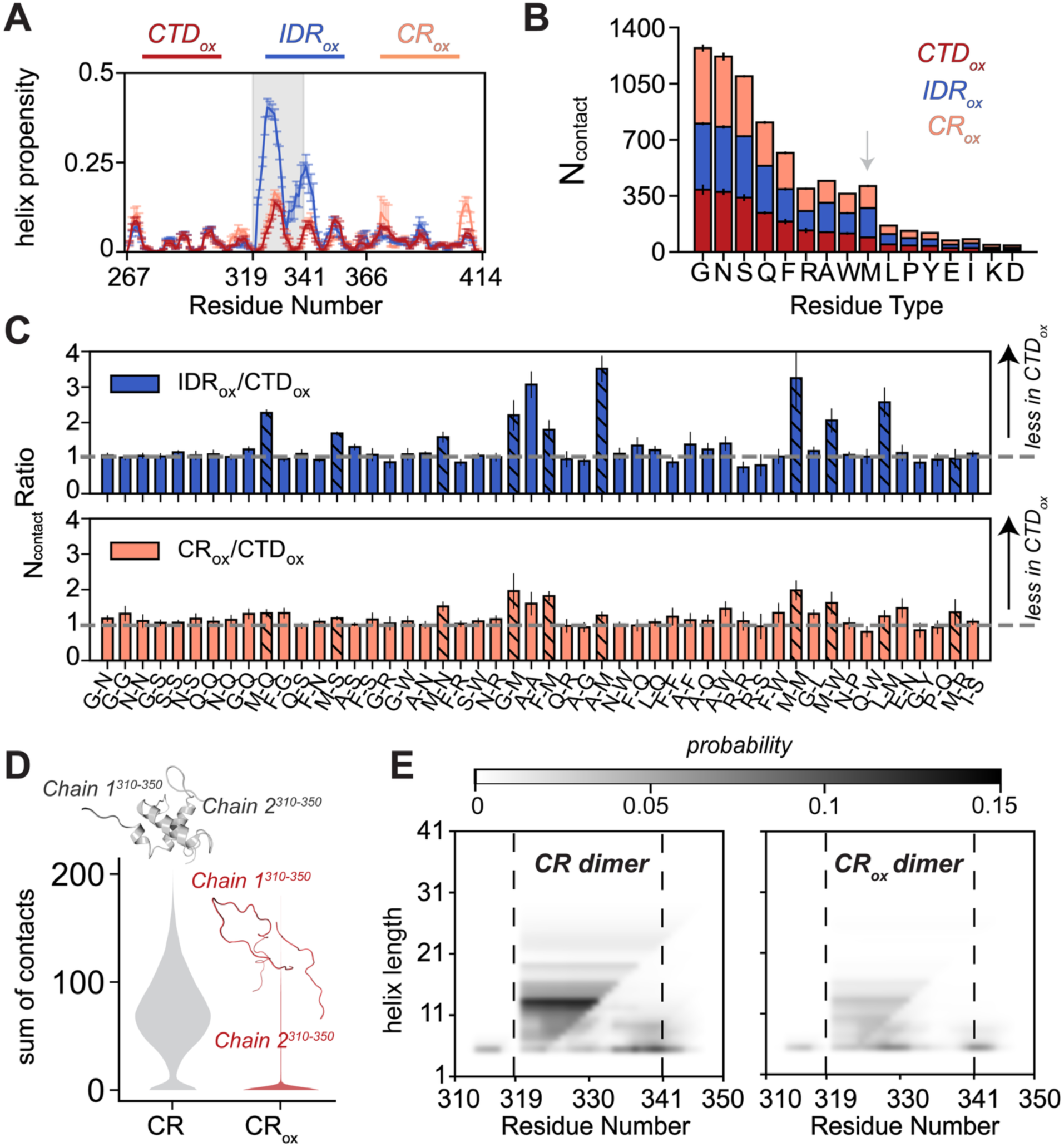
Methionine oxidation in distinct regions of TDP-43 CTD differentially affects structural properties and interaction profiles. **(A)** Helical propensity of CTD. CR methionine oxidation (CR_ox_) dramatically reduces helicity to levels comparable to complete oxidation, while IDR methionine oxidation (IDR_ox_) minimally affects helical structure relative to unoxidized CTD (gray), demonstrating the critical role of CR methionines in maintaining structural integrity. **(B)** Number of contacts formed by each residue type is shown for oxidative variants. **(C)** Quantification of contacts between specific residue types. Ratio is calculated by dividing number of contacts for each residue pair in IDR_ox_ and CR_ox_ with CTD_ox_. **(D)** CR-CR dimer interface stability. AlphaFold v2.3 dimer models were subjected to 250 ns MD simulations with and without methionine oxidation. Distribution quantifies CR-CR contacts across all simulations, showing significant disruption of intermolecular interactions upon oxidation. Inset: Representative structures from simulations showing intact (CR) and disrupted (CR_ox_) dimer interfaces. **(E)** Averaged helix length propensity across the CR region (residues A310-S350) for CR (left) and CR_ox_ (right) simulations. Helix length was computed for both chains across trajectories and averaged. The heatmap reveals consistent reduction in helical content throughout the CR sequence upon methionine oxidation.

Further investigation of the CTD intramolecular (pairwise) interaction profile revealed that both oxidation patterns altered the intramolecular interaction network in a manner similar to complete oxidation (**Figure 3E, 5B, 5C**), resulting in fewer intramolecular interactions compared to the unoxidized ensemble (**Figure S13**). Position-specific summation of pairwise contacts showed pronounced reduction only in the C-terminal portion of IDR1 for the IDR_ox_ ensemble, whereas CR_ox_ showed contact disruption in both CR and Q/N regions. Analysis of specific residue-type pairs revealed that Met interactions were disrupted to different extents in the two variants relative to completely oxidized CTD (**Figure 5B-C, S14 and S15**). For IDR_ox_, with the exception of M-R (contact ratio∼1), Met interactions with polar (Q, N, S), aliphatic (A, L, M), and aromatic (F, W) residues were less disrupted than those in CTD_ox_, as indicated by contact ratios (IDR_ox_/CTD_ox_) exceeding 1.5. In contrast, CR_ox_ showed disruption comparable to CTD_ox_, with contact ratios approaching 1, consistent with the greater chain expansion observed for both CR_ox_ and CTD_ox_ ensembles.

Furthermore, examination of region-level interactions revealed that intra-region interactions outside the CR remained largely similar to unoxidized CTD in both variants (**Figure S16**). However, the inter-region interaction network was dramatically altered in both cases through different mechanisms. In CR_ox_, the disruption of helical structure directly impaired CR-CR interactions, while also negatively impacting majority of the CR interactions with other regions (**Figure S16**). Despite preserving CR helicity, the oxidation of CR-flanking Met residues in IDR_ox_ still reduced inter-region interactions involving IDR1-Q/N, IDR1-IDR2, and CR-IDR2 compared to unoxidized CTD (**Figure S16**). Ultimately, oxidation of Met residues, whether in CR or IDR, significantly disrupted long-range interactions between N- and C-terminal domains, leading to more expanded ensembles.

The observed reduction in helicity and CR-CR interactions upon Met oxidation in the conserved region prompted us to investigate how these molecular changes affect helix-helix interactions, which represent a critical initial step in the self-assembly process. We generated CR dimer models (residues 310-350) using AlphaFold v2.3^25^ and performed AAMD simulations (250 ns x 18 models) comparing unoxidized to oxidized (CR_ox_) dimers (**Figure S17**). These simulations revealed that Met oxidation triggered rapid dissociation of CR dimers, with most models showing significant loss of CR-CR contacts during early simulation stages (**Figure 5D and S18**). Importantly, this dimer dissociation strongly correlated with loss of CR helical structure (**Figure 5E**).

In summary, our results demonstrate that Met oxidation in different regions of the TDP-43 CTD disrupts self-assembly and phase separation through distinct molecular mechanisms. Oxidation of IDR Met residues primarily alters the physicochemical properties of these flanking regions, weakening the intermolecular interactions necessary for condensate formation while largely preserving helical structure. In contrast, Met oxidation within the CR directly compromises helical stability, causing rapid dimer dissociation and preventing the initial stages of self-assembly. These region-specific effects collectively highlight the balance of interactions required for phase separation, demonstrating how oxidative modifications may exert functional consequences through alternative molecular pathways.

## CONCLUSIONS

Recently, a link has been suggested between oxidation and TDP-43’s disease-associated dysfunction – Cys oxidation can promote TDP-43 aggregation through inter- and intra-molecular disulfide bridges and other oxidized states and cross-links^65–67^. Here we now show the mechanism by which reversible Met oxidation may play a distinct regulatory role in TDP-43 function. The functional significance of Met residues is highlighted by our previous work establishing them as critical contributors to TDP-43 phase separation^54^. Furthermore, the disease-associated M337V mutation affects cellular redox responses by altering interactions with heterogeneous nuclear ribonucleoprotein K (hnRNP K) under stress conditions^68,69^, suggesting Met residues may serve as redox-sensitive modulators of protein function rather than simply driving pathological aggregation. As TDP-43 CTD harbors ten Met residues positioned throughout both the CR and flanking regions, their region-specific roles in oxidative stress response may be important yet have remained elusive. In this study, we demonstrate that oxidation of Met residues differentially impacts TDP-43 CTD based on their location: CR oxidation disrupts helical structure essential for self-assembly, while oxidation in the flanking regions alters the interaction landscape, both contributing to impaired phase separation.

Our experimental analyses revealed that all ten TDP-43 CTD Met residues are susceptible to oxidation, leading to profound changes in the CTD’s conformational ensemble. Our complementary AAMD simulations provided mechanistic insights into these changes, showing that reduced hydrophobicity upon oxidation triggers a dramatic reorganization of the CTD interaction network with reduced intramolecular contacts involving polar and hydrophobic residues. This reorganization manifests as conformational expansion at the monomer-level through the weakening of long-range interactions. Both our NMR data and computational analyses demonstrated that these changes result in substantial disruption of CR helicity. The combination of disrupted helicity and long-range interactions explains the impaired phase separation we observed by DIC microscopy, as these changes negatively impacted the helix-helix interactions, a critical step in higher-order self-assembly (**Figure 6**).

**Figure 6.**
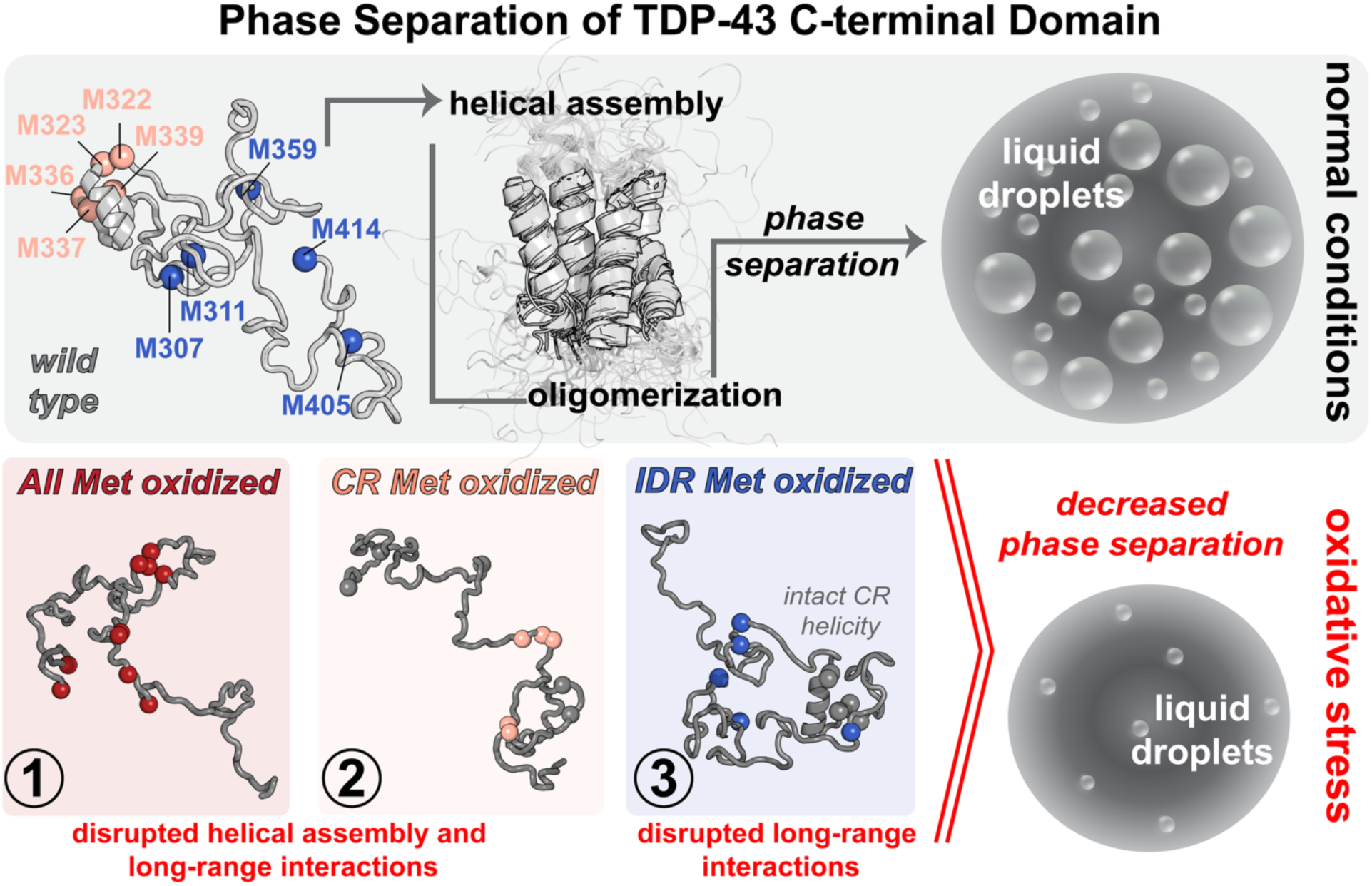
Methionine oxidation decreases phase separation propensity of TDP-43 CTD. Schematic illustrating the effects of methionine oxidation on TDP-43 CTD structure and phase behavior. Under normal conditions, TDP-43 CTD helical assembly leads to oligomerization triggering phase separation and droplet formation. Under oxidative stress, methionine oxidation disrupts phase separation through distinct mechanisms: (1) oxidation of all methionines or (2) methionines at conserved region (CR) alone disrupts both the formation of helical assembly and long-range interactions involving polar and hydrophobic residues, while (3) oxidation of methionines at IDR1 preserves CR helicity but prevents stable helical assembly by rewiring protein-protein interaction network. All scenarios result in decreased phase separation propensity.

To investigate position-specific effects of Met oxidation on CTD phase separation, we created variants where Met residues were systematically removed or relocated from either the IDRs or in the helical CR. Regardless of Met positioning, oxidation consistently disrupts phase separation, yet does so via two distinct but complementary mechanisms: in the flanking IDR regions, Met oxidation weakens multivalent interactions required for phase separation, while in the CR region, oxidation directly destabilizes helical conformations and helical interactions. Ultimately, both variants exhibit conformational expansion through loss of critical long-range contacts. To specifically examine the effect of oxidation on intermolecular association, we further conducted AAMD simulations starting from an ensemble of AlphaFold-predicted dimer models with and without CR Met oxidation. These simulations showed rapid dissociation of the CR-CR interface upon oxidation, directly demonstrating how oxidized CR Met residues disrupt the helix-helix interactions that drive phase separation. Together, our findings suggest that disruption of CR helicity and/or long-range interactions is sufficient to prevent phase separation, highlighting the delicate balance of interactions maintaining CTD’s functional state (**Figure 6**).

In conclusion, our study elucidates how Met oxidation systematically disrupts the structure and phase separation of the TDP-43 CTD, offering molecular-level insights into protein phase behavior under oxidative stress conditions. Further, we reveal the dual role of Met residues in CTD: as important contributors to CR helical stability and as redox-sensitive modulators of phase separation. Consequently, our findings provide molecular insights into how oxidative stress may affect TDP-43 phase behavior and offer a framework for understanding mechanisms of proteinopathy-related dysfunction.

## Acknowledgements

This work was supported by **NINDS and NIA R01NS116176**. All-atom MD simulations were conducted with the advanced computing resources provided by Texas A&M High Performance Research Computing.

## Author contributions

B.O, R.Z.P., and R.V. contributed equally to this work. R.Z.P., R.V., J.S., and N.L.F. designed and analyzed the biochemical and biophysical experiments, and R.Z.P., R.V., and J.S performed the experiments. B.O., P.M., and J.M. designed, performed and analyzed data for MD simulations. B.O., P.M., R.Z.P., R.V., J.M. and N.L.F wrote the manuscript with contributions from all authors.

## Conflict of Interest Statement

The authors declare no other conflicts of interest.

## MATERIALS AND METHODS

### Experimental Procedures

#### Recombinant protein expression and purification

The TDP-43 variant plasmids were generated through by codon-optimization and gene synthesis sequences and the protein products were expressed within the pJ411 bacterial expression vector. The expression process was conducted in BL21 Star (DE3) *Escherichia coli* cells (Life Technologies), cultivated in LB or, for NMR experiments, M9 minimal media supplemented with ^15^NH_4_Cl. The expression and purification methodology followed a slightly adapted protocol from Mohanty et al PNAS 2023^54^ as follows.

Bacterial cultures were induced at an optical density (OD) of 0.8 with 1 mM IPTG, followed by a 4-hour incubation at 37°C and 220 rpm. Subsequently, cells were harvested through centrifugation (6000 rpm, 15 minutes, 4°C), and the resulting cell pellets were resuspended in a 20 mL buffer (20 mM Tris, 500 mM NaCl, 10 mM imidazole, pH 8.0). Cells were then lysed by using an ultrasonic cell disruptor, followed by centrifugation (15,000 g, 1 hour, 4°C). The insoluble fraction, containing inclusion bodies, were resuspended in a 40 mL solubilizing buffer (8M urea, 20 mM Tris, 500 mM NaCl, 10 mM imidazole, pH 8.0) and centrifuged (25,000 rpm, 1 hour, 19°C).

The supernatant was then filtered using a 0.45 μm syringe filter, and protein purification was carried out using a 5 mL Histrap HP column with a gradient of 10 to 500 mM imidazole added to the solubilizing buffer. The purified protein fractions underwent desalting with a HiPrep 26/60 Desalting Column into TEV cleavage buffer (20 mM Tris, 500 mM GdnHCl, pH 8.0) and were subjected to TEV cleavage overnight at room temperature. Post-cleavage, solid urea and NaCl were added to achieve a concentration of approximately 8 M urea and 500 mM NaCl. The redissolved protein was applied to the Histrap HP column for the removal of the histidine tag and histidine-tagged TEV protease. The cleaved protein fractions were concentrated, buffer-exchanged into a storage buffer (20 mM MES, 8 M urea, pH 6.1), flash-frozen, and stored at −80°C for future use.

#### Oxidation of Protein Samples

Protein samples in storage buffer were quickly thawed and split into two samples of equal volume. The oxidation reaction was performed by addition 3% w/v aqueous H_2_O_2_ to one sample to achieve a final H_2_O_2_ concentration of 1% w/v. A equivalent volume of water was added to the other sample to act as unoxidized control. After 30 min the reaction was slowed by 10-fold dilution of both samples in storage buffer. Both the oxidized sample and unoxidized sample were then buffer-exchanged back into storage buffer, concentrated to a protein concentration of 2 mM, flash-frozen, and stored at -80°C for future use.

#### Mass spectrometry

LC/MS analyses were conducted on an Agilent 1290 Infinity II LC coupled with an Agilent 6520 TOF system equipped with a dual ESI source and a Phenomenex Jupiter C4 5 µm 300 A column (150 × 2.0 mm). The mobile phases consisted of 0.1% formic acid in water (A) and 0.1% formic acid in acetonitrile (B). A gradient of 17 min of 5–90 % B at the flow rate of 0.4 ml/min was used. The gradient program was as follows: 5%-60% B (0-7 min), 60%-90% B (7-12 min), and 90% B (12-17 min). 1 µl sample at a concentration of 20 µM was injected. MS spectra were acquired in positive ionization mode over a mass range of m/z 100–10000 with an acquisition time of 1.0 s per spectrum. The ESI source was operated with the following parameters: capillary voltage at 3500 V, nebulizer 35 psi, fragmentor voltage 175V, drying gas (nitrogen) flow rate at 11 l/min, and drying gas temperature at 325 °C. The mass method was segmented. The eluent of the first segment (0-2.5 min) was diverted to waste. Instrument control and data analysis were carried out using MassHunter 10.0 software. The deconvolution was conducted using MassHunter BioConfirm 10.0 software.

#### Microscopy

Adhering to the manufacturer’s guidelines, desalting of the protein samples was performed employing 0.5 mL Zeba spin desalting columns purchased from Thermo Fisher Scientific. In this desalting procedure, the concentrated protein stocks were first diluted from 8 M urea to 1 M urea with the MES sample buffer (20 mM MES, pH 6.1), applied to the desalting column, spun, and collected, after which the desalted protein samples were diluted to 80 μM with either 20 mM MES, pH 6.1 or 20 mM MES, pH 6.1 supplemented with 150 mM NaCl. After gentle mixing, the samples were observed through DIC microscopy, using a Nikon Ti2-E Fluorescence Microscope equipped with a 40x objective. For image acquisition, 10 μL of each sample was deposited onto a coverslip. Images were processed using Fiji software.

#### NMR sample preparation and spectroscopy

Nuclear magnetic resonance (NMR) spectroscopy was carried out using Bruker Avance spectrometers at 850 MHz ^1^H Larmor frequencies, equipped with HCN TCI z-gradient cryoprobes and maintained at sample temperature of 298 K. NMR samples, were prepared by dilution of 2 mM protein stocks to the desired concentration with 20 mM MES pH 6.1 with 5.23% added ^2^H_2_O with 5% ^2^H_2_O (for NMR locking) sample buffer to achieve a final 5% ^2^H_2_O and 360 mM residual urea concentration for all samples regardless of protein concentration (i.e. a 500 μl sample consistent of 22.5 μl of protein stock in 8 M urea storage buffer with additional 8 M urea storage buffer which was then added to 477.5 μl of 20 mM MES pH 6.1 5.23% ^2^H_2_O). NMR data was processed using NMRPipe^70^ and analyzed using NMRFAM SPARKY^71^.

Backbone assignments for the WT, 5M→A^IDR^, and 5M→A^CR^ were transferred from the Biological Magnetic Resonance Bank (BMRB) records (BMRB codes 26823, 52062, and 52060 respectively). Assignment of 5M→A^CR^+ 5A→M^IDR^ CR region was achieved by overlay with 5M→A^CR^ assignments. Backbone assignment of oxidized variants was achieved through standard triple resonance experiments (HNCACB, HNCO, HNCA, HNN and CBCACONH) using experimental parameters described previously^24^. ^1^H-^13^C HSQCs were collected of samples with 512 and 4096 points in the indirect ^13^C and direct ^1^H dimensions with acquisition times of 15 ms and 200 ms with sweep widths of 76.0 ppm and 12.0 ppm centered around 38.0 ppm and 4.7 ppm, respectively. (ΔδC_α_-δC_β_) were calculated as follows: ΔδC_α_-δC_β_ = (δC_αexp_-δC_αref_) -(δC_βexp_-δC_βref_), where δC_αexp_ and δC_βexp_ are the experimentally measured C_α_ and C_β_ chemical shifts, and δC_αref_ and δC_βref_ are the calculated Cα and Cβ random coil reference chemical shifts computed using the Poulsen IDP/IUP random coil chemical shifts calculator using default parameters and the experimental temperature (http://spin.niddk.nih.gov/bax/nmrserver/Poulsen_rc_CS/)^72,73^ respectively. ^1^H-^15^N HSQCs were collected of samples with 200* and 2048* complex pairs in the indirect ^15^N and direct ^1^H dimensions with acquisition times of 105 ms and 200 ms and sweep widths of 22.0 ppm and 12.0 ppm centered around 116.5 ppm and 4.7 ppm, respectively. The ^1^H and ^15^N chemical shifts (Δδ^1^H and Δδ^15^N) were calculated as the difference in the chemical shift of resonances at a sample concentration of 90 μM minus the chemical shift at a sample concentration of 20 μM. The combined chemical shift (Δδ^1^H-^15^N) was calculated as follows: 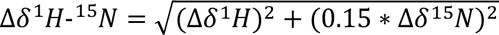. ^15^N *R_2_* of 5M→A^CR^ and 5M→A^CR^_Ox_ was measured at 850 MHz ^1^H using a standard pulse sequence (hsqct2etf3gpsitc3d from Bruker Topspin 3.2) acquired with 128* and 1536* complex pairs in the indirect ^15^N and direct ^1^H dimensions with acquisition times of 78 ms and 150 ms and sweep widths of 19.0 ppm and 12.0 ppm centered around 116.5 ppm and 4.7 ppm, respectively. ^15^N R2 experiments had an interscan delay of 2.5 s, a Carr-Purcell-Meiboom-Gill (CPMG) field of 556 Hz, and total R_2_ relaxation CMPG loop-lengths of 16.3 ms, 32.6 ms, 48.9 ms, 65.2 ms, 163.0 ms, 195.6 ms, and 244.5 ms.

### Computational Procedures

#### Molecular Dynamic (MD) Simulations of Methionine Oxidation

Initial configurations for triplicate MD simulations of the TDP-43 CTD were obtained by performing k-means clustering on the CTD ensemble^54^ using the “sklearn” module in Python. The centroids of the three most populated clusters were submitted to the Vienna-PTM 2.0^74^ webserver to introduce oxidation. Since there is no preferential formation of one diastereomer of Met sulfoxide in proteins^75^, we modified Methionine to Methionine (R) sulfoxide using parameters generated by Irani et al. (2013)^76^. To generate CTD_ox_, we modified all Met residues except M414 at the C-terminus; for CR_ox_, we modified M322/323/336/337/339; and for IDR_ox_, we modified M307/311/359/405. M414 at the C-terminus was excluded as the force field parameters for a C-terminal methionine sulfoxide were not readily available.

The structures were prepared for simulations in GROMACS 2021^77^ using the Amber99sb-stq force field^78^ and the TIP4P/2005s water model^79^. The simulation box was a truncated octahedron with a length of 12.5 nm, and counter ions were added to maintain electroneutrality and a salt concentration of 0.10 M NaCl. After minimization, the GROMACS files (“top” and “gro”) were converted to AMBER files (“parm7” and “rst7”) using ParmED^80^. During the conversion, hydrogen mass repartitioning^81^ to 1.5 amu was performed to enable a timestep of 4 fs.

Following system preparation, production runs were carried out in Amber22^82^ at 1 bar and 300 K, after an equilibration process. The energy minimization was conducted using the steepest descent and conjugate gradient algorithms, with all non-hydrogen atoms of the protein restrained by a 5 kcal/(mol Å) force constant. The energy-minimized structures were heated for 1 ns with a 2 fs time step, with the temperature linearly increasing from 0 to 300 K for 0.2 ns and then maintained at 300 K for the remainder. During this stage, the force constant was reduced to 0.5 kcal/(mol Å) and subsequently removed for an additional 5 ns NVT equilibration. The time step was increased to 4 fs, followed by another 10 ns NVT equilibration. A density equilibration was then performed under an isothermal-isobaric ensemble (NPT) using the Berendsen barostat^83^ for 10 ns. The production run was conducted using the Monte Carlo barostat^84^ with an isotropic coupling of 1.0 ps at a pressure of 1 bar. Temperature control was achieved using Langevin dynamics with a 1.0 ps^−1^ friction coefficient, and bonds involving hydrogen atoms were constrained using SHAKE^85^. A nonbonded cutoff of 0.9 nm was applied for short-range nonbonded interactions, and long-range electrostatic interactions were treated using the particle mesh Ewald method^86^. Triplicate runs of 5 µs each were performed.

To confirm that the overall length of our individual trajectories (5 μs each) were of sufficient length for conformational exploration beyond the initial state, we calculated radius of gyration (R_g_) (**Figure S2A and S9A**) which is a well-established measure of chain global dimensions. Further, we estimated the rate of chain relaxation by calculating the time autocorrelation function of R_g_ over each of the trajectories (**Figure S2B and S9B**). The average correlation times were 121, 187, 108, and 146 ns for CTD, CTD_ox_, IDR_ox_, CR_ox_ respectively, which is nearly 30-fold smaller than the trajectory duration, thereby confirming that both ensembles rapidly deviate from their initial configuration.

Further, the adoption of multi-microsecond simulation replicas can improve sampling efficiency and enable a wider exploration of the conformational landscape compared to the individual replicas^87–89^, as indicated by the distinct (non-overlapping) peaks in the R_g_ distribution observed across the replicas of variants (**Figure S2C, and S9C**).

#### Molecular Dynamic (MD) Simulations of CR Dimers

Initial configurations (eighteen models) for dimer simulations were generated using AlphaFold v2.3 in our previous study^90^. For oxidation simulations, all CR Met residues in both chains were subjected to the MetOx modification and prepared for simulations in GROMACS as described above. Both unoxidized and modified dimer structures underwent equilibration followed by 250 ns production runs in Amber22 under the NPT ensemble.

The equilibration process consisted of several sequential steps: (1) energy minimization with restraints (force constant 5 kcal/mol·Å²); (2) 1 ns heating from 0 K to 300 K with restraints (force constant 5 kcal/mol·Å²); (3) seven stages of gradual restraint release under NVT ensemble totaling 7 ns, with force constants decreasing stepwise (2.5, 1, 0.5, 0.25, 0.10, 0.05, 0.01 kcal/mol·Å²); (4) 2 ns NVT equilibration without restraints; (5) 5 ns NVT equilibration with increased timestep (4 fs); and (6) density equilibration consisting of two 5 ns simulations, first with Berendsen barostat followed by Monte Carlo barostat. All other simulation parameters matched those used in the single chain simulations.

#### Structural and Contact Analysis

GROMACS utility tools were used for Radius of Gyration (Rg). Time evolution of secondary structure analysis is conducted in GROMACS according to Kabsch and Sander’s DSSP algorithm^91^. All contact analysis were performed using MDAnalysis “neighbor search library” module^92,93^. Protein-protein contacts were characterized by a 4.5 Å distance cutoff between heavy atoms. A contact was considered formed if a residue had one or more heavy atoms within 4.5 Å of another residue’s atoms. Molecular visualizations were rendered using PyMOL^94^.

## Notes

### Competing Interest Statement

The authors have declared no competing interest.

