## Supplementary Information for "Site-specific methionine oxidation alters structure and phase separation of TDP-43 C-terminal domain"

[1]Artie McFerrin Department of Chemical Engineering, Texas A&M College of Engineering, College Station, Texas; [2]Therapeutic Sciences Graduate Program, Brown University, Providence, Rhode Island; [3]Department of Molecular Biology, Cell Biology & Biochemistry, Brown University, Providence, Rhode Island; [4]Department of Chemistry, Texas A&M University, College Station, Texas; [5]Interdisciplinary Graduate Program in Genetics and Genomics, Texas A&M University, College Station, Texas

* B.O, R.Z.P., and R.V. contributed equally to this work.

[#] Correspondence should be addressed to:

J.M. and N.F.

**A**

```
Human      RQLERSGRFGGNPGGFGNQGGFGNSRGGGAGLGNNQGSNMG--GGMNFGAFSINPAMMAAAQAALQSSWGMMGMLA 341
Mouse      RQLERSGRFGGNPGGFGNQGGFGNSRGGGAGLGNNQGGNMG--GGMNFGAFSINPAMMAAAQAALQSSWGMMGMLA 341
Chicken    RQLERGGRFGGNPGGFGNQGGFGNSRGGGGGLGNNQGSNMG--GGMNFGAFSINPAMMAAAQAALQSSWGMMGMLA 341
Frog       RQLERGGRFPGPS--FG-NQGYPNSRPSSGALGNNQGGNMGGGGGMNFGAFSINPAMMAAAQAALQSSWGMMGMLA 349
Zebrafish  QMMERAGRFGNGFGGQGFAGSRSNMGGGGGGS----SS-----SLGNFGNFNLNPAMMAAAQAALQSSWGMMGMLA 341
           : :**.*** .     *   .  *   ....     ..     .  *** *.:***********************

Human      SQQNQSGPSGNNQNQGNMQREPNQAFGSGNNSYSGSNSGAAIGWGSASNAGS-GSGFNGGFGSSMDSKSSGWGM  414
Mouse      SQQNQSGPSGNNQSQGSMQREPNQAFGSGNNSYSGSNSGAPLGWGSASNAGS-GSGFNGGFGSSMDSKSSGWGM  414
Chicken    SQQNQSGPSGNNQPQGNMQREQNQGFSSGNNSYGGSNSGAAIGWGSASNAGS-SSGFNGGFGSSMDSKSSGWGM  414
Frog       SQQNQSGPQGSNQGQGNQQRDQPQSFGSNN-SYGS--NSGAIGWGSP-NAGS-GSGFNGGFSSSMESKSSGWGM  418
Zebrafish  -QQNQSGTSGTSTSGTSSSRDQAQTYSSANSNYGS--SSAALGWGTGSNSGAASAGFNSSFGSSMESKSSGWGM  412
           .****** .*..    . .*:  * :.* * .*..  ... :***:  *:*: .:***..*.***:********
```

**B**

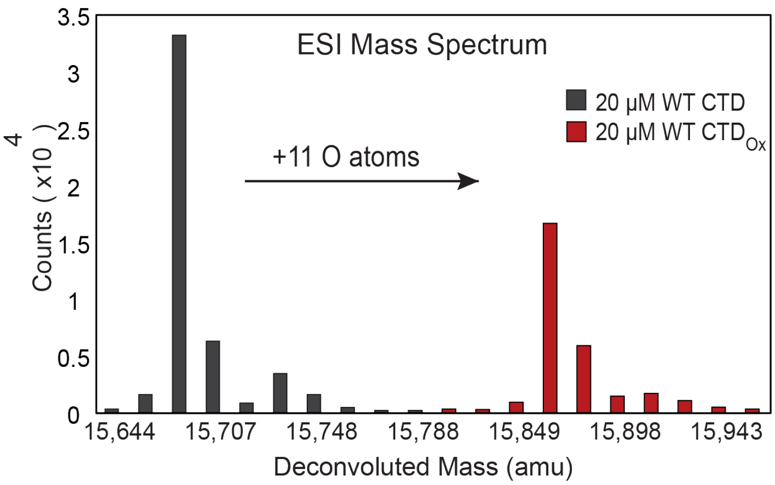

**Figure S1. TDP-43 CTD methionine residues are highly conserved across vertebrates and are targets of oxidation**. (A) Multiple sequence alignment of TDP-43 CTD from vertebrate model organisms including human, mouse, chicken, frog, and zebrafish demonstrating the position of methionine residues is highly conserved across vertebrate organisms. (B) Mass spectrum of WT TDP-43 CTD before and after oxidation with 1% v/v $H_2O_2$ reveals a shift in the mass of the predominant species corresponding to the addition of an oxygen atoms to CTD methionine residues following the conversion of thioether groups to sulfoxide.

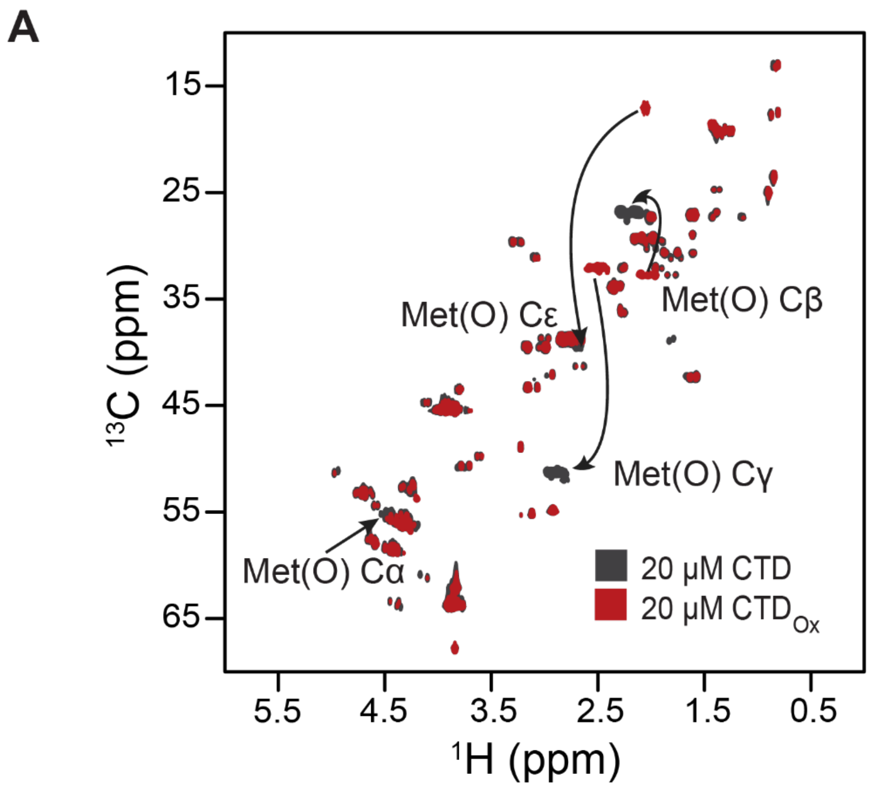

**Figure S2. NMR spectra confirms oxidation of all methionine residues. (A)** Overlay of 1H-13C HSQC spectra of 20 μM unoxidized WT CTD (grey) and 20 μM oxidized WT CTD (red) demonstrating that methionine sidechain carbons undergo dramatic CSPs following oxidation of the sulfur-containing thioether group.

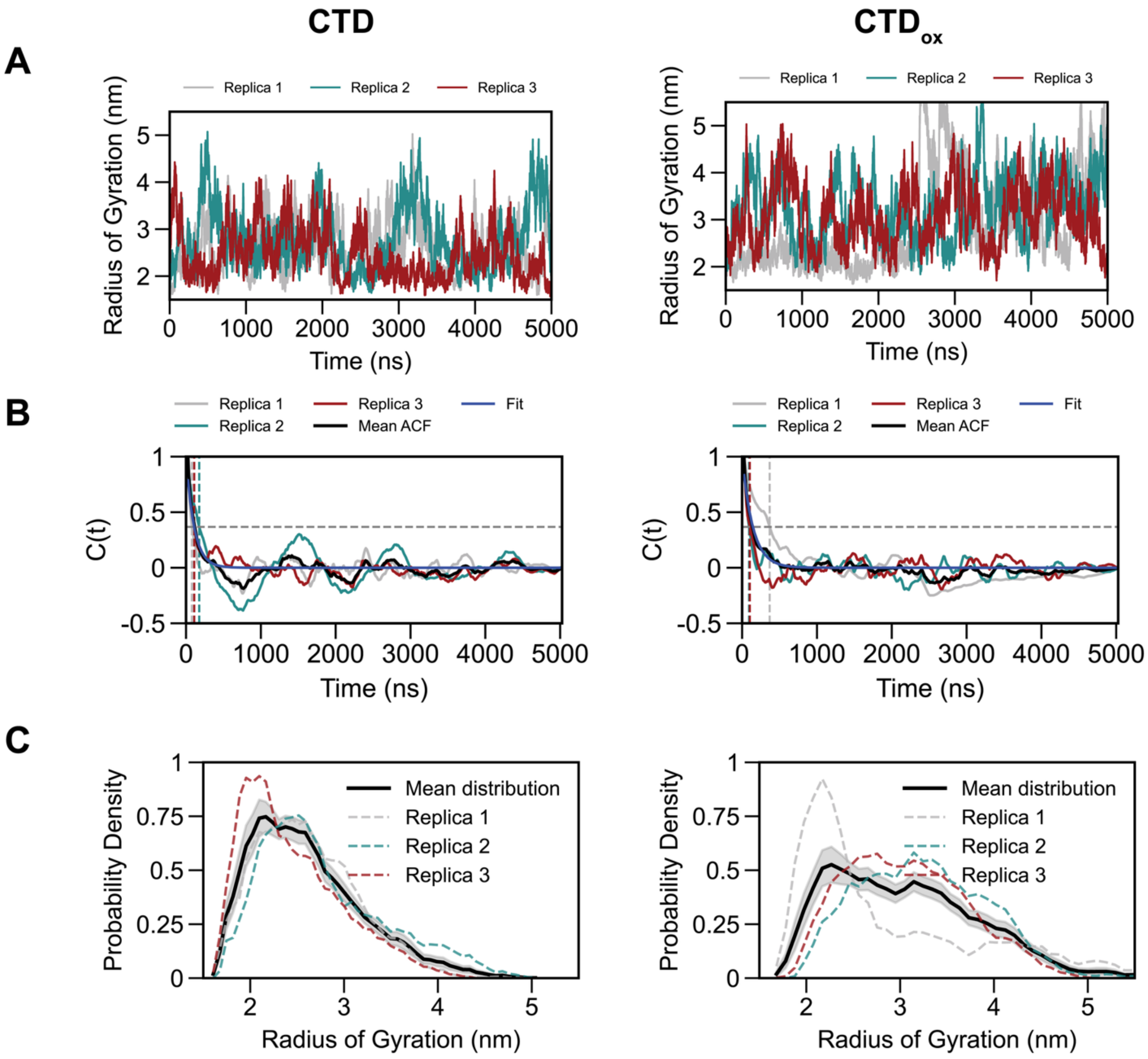

**Figure S3. Assessing the extent of conformational sampling across multi-replica trajectories of CTD and CTD$_{ox}$. (A)** Rg variation as a function of time over three independent replica trajectories of CTD and CTD$_{ox}$. **(B)** The plots show the time evolution of the radius of gyration autocorrelation function C(t) for each variant with data from three independent replicas. C(t) was calculated from the backbone radius of gyration (Rg) for each individual trajectory. The vertical dotted lines indicate where the autocorrelation function crosses 1/e (≈ 0.3679). Based on these crossing times, we conservatively selected 500 ns as the equilibration time for all systems. After this period, the autocorrelation functions consistently fluctuate around zero, indicating decorrelated sampling suitable for subsequent analysis. **(C)** Probability distributions of radius of gyration (Rg) with data from three independent replicas (shown in gray, blue, and red dashed lines) and their mean distribution (solid black line).

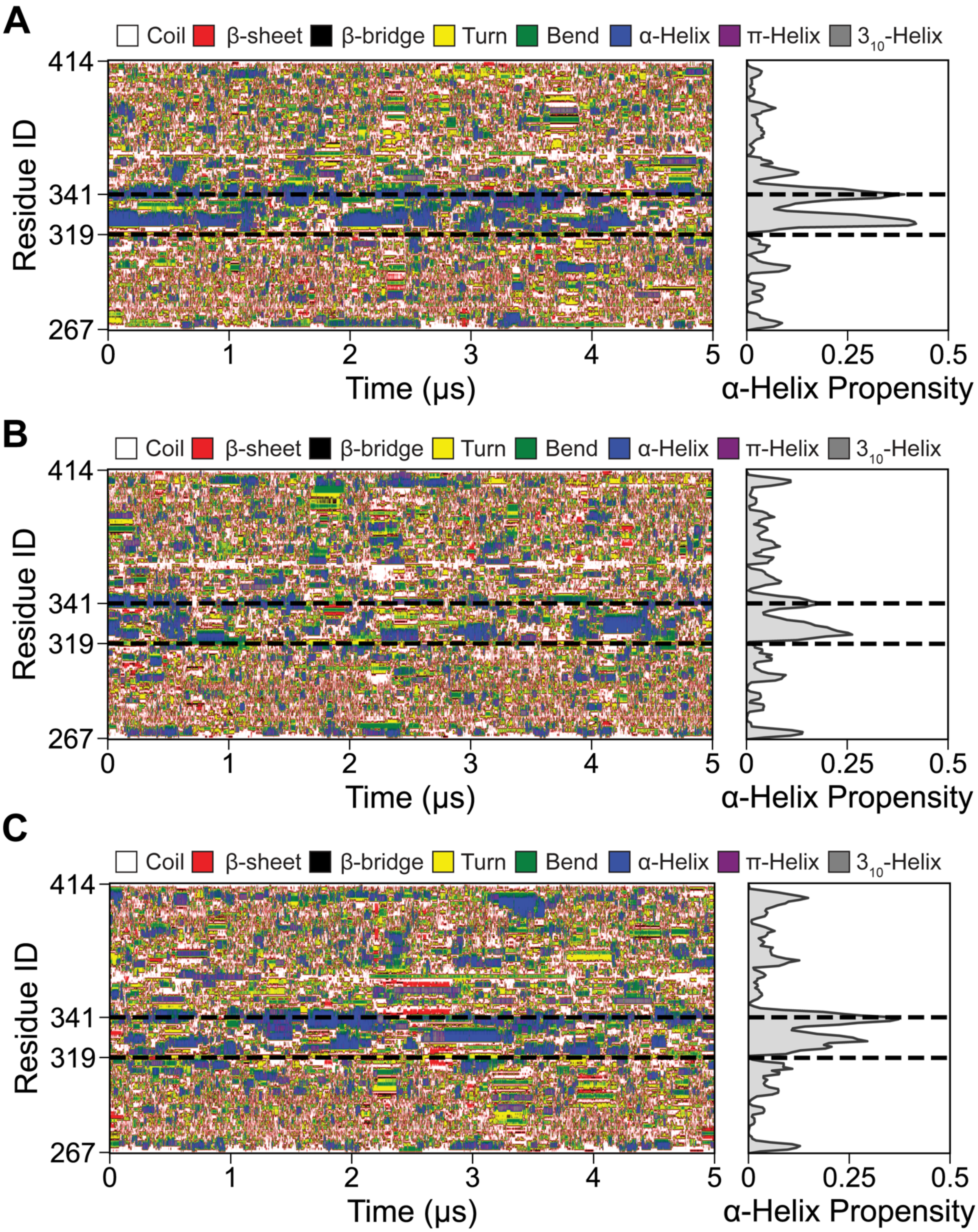

**Figure S4. Time evolution of secondary-structure changes in CTD**. **(A)** Replica 1, **(B)** Replica 2, and **(C)** Replica 3 showing residue-level secondary structure transitions over 5 μs.

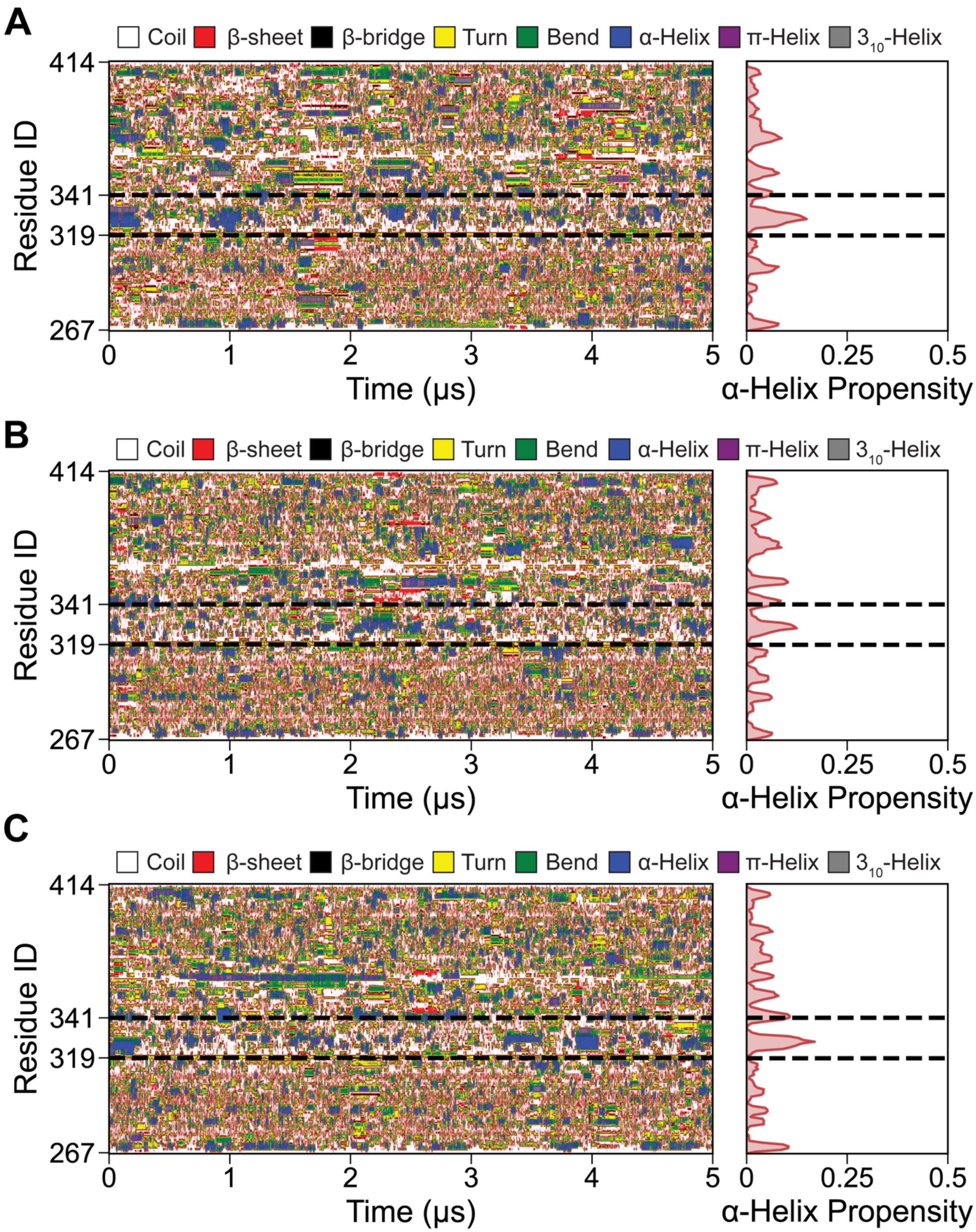

**Figure S5. Time evolution of secondary-structure changes in $CTD_{ox}$ (A)** Replica 1, **(B)** Replica 2, and **(C)** Replica 3 showing residue-level secondary structure transitions over 5 μs.

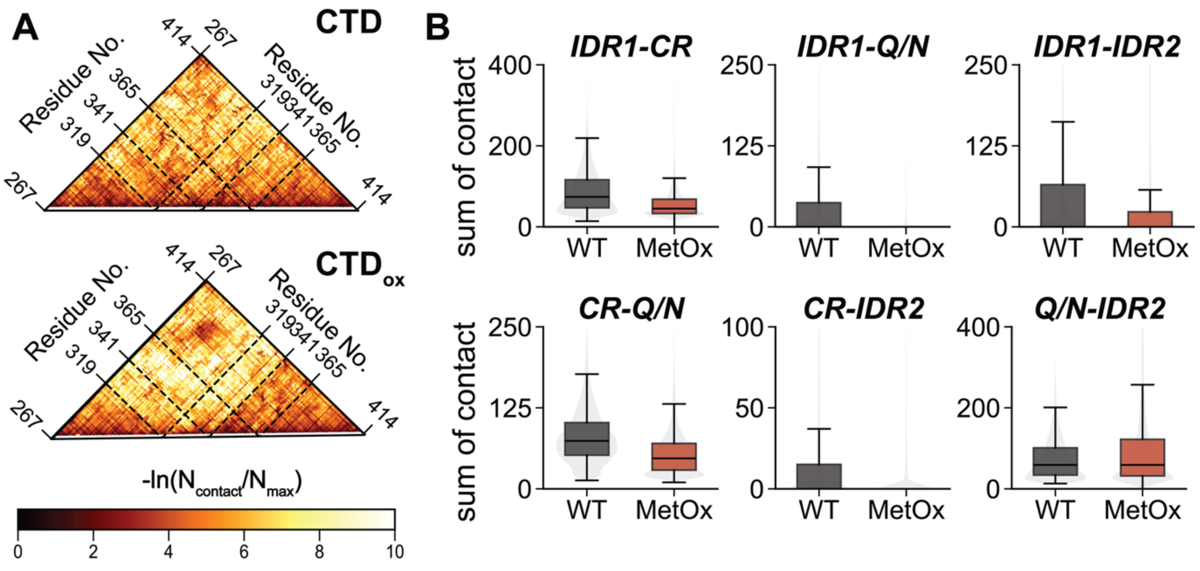

**Figure S6. Analysis of TDP-43 CTD intra-protein contact networks under oxidation conditions**. **(A)** Contact maps revealing intraprotein interactions in CTD (top) and CTD$_{ox}$ (bottom) conditions. Contact frequencies are represented as $-\ln(N_{contact}/N_{max}^{CTD})$, where darker red indicates higher contact probability. Region boundaries (IDR1, CR, Q/N, IDR2) are marked with dashed lines. **(B)** Effect of oxidation on long-range interactions within CTD. For each frame, all heavy atom contacts (4.5 Å cutoff) between protein residue pairs were calculated and grouped by region: IDR1 (residues 267-318), CR (residues 319-341), Q/N (residues 342-365), and IDR2 (residues 366-414). Both intra-region (e.g., IDR1-IDR1) and inter-region (e.g., IDR1-CR) interactions are shown. Box plots display medians (solid black lines) and means (dashed black lines), with boxes extending from first (Q1) to third (Q3) quartiles. Points beyond 1.5 times the interquartile range from either Q1 or Q3 are plotted as outliers. Gray violin plots in the background reveal the full probability distribution of contact frequencies across all analyzed frames, highlighting differences in interaction patterns between CTD (gray) and CTD$_{ox}$ (red) proteins.

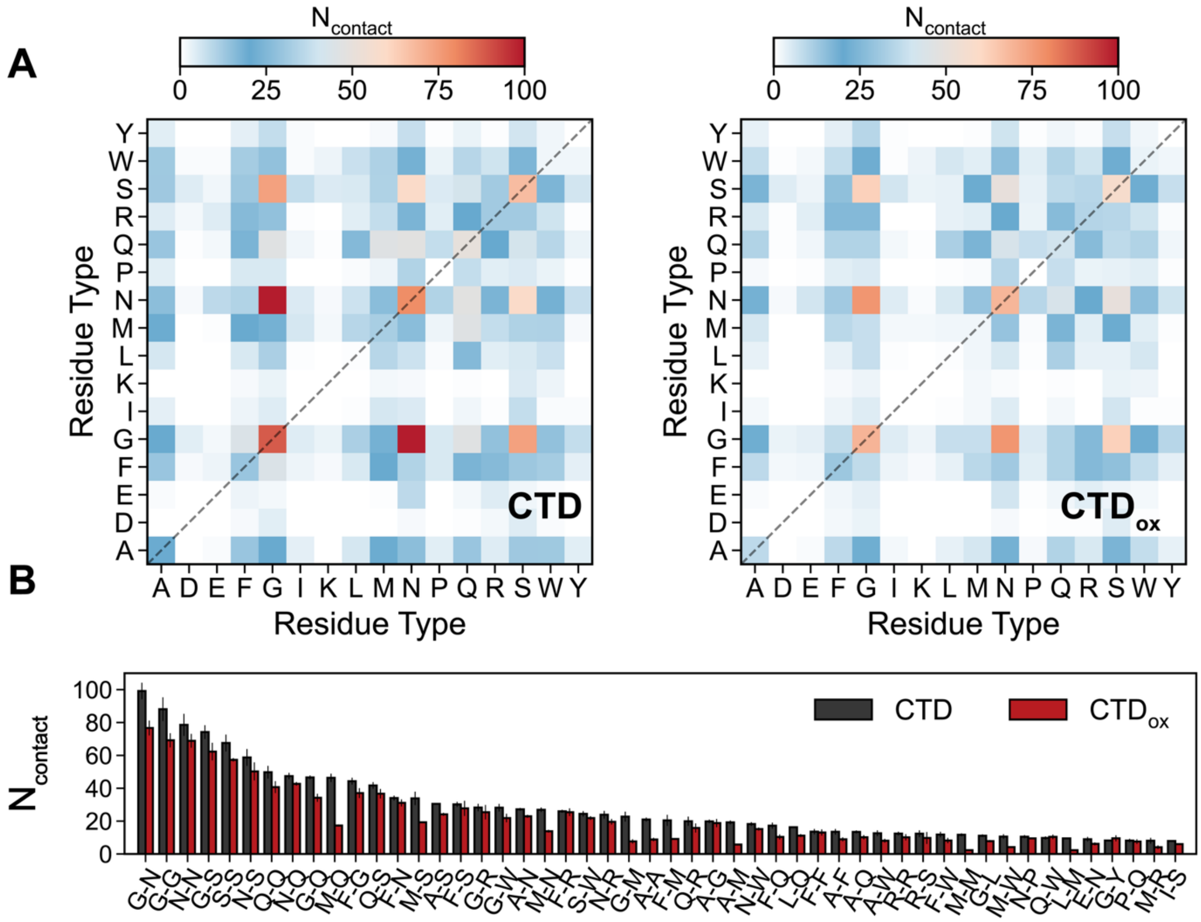

**Figure S7. Comprehensive Residue Type Interaction Analysis** (A) Interaction heatmap displaying contact frequencies between different amino acid types for CTD and upon Met oxidation (CTD$_{ox}$) conditions. The color scale ranges from blue (low contact) to red (high contact), with the intensity indicating the number of interactions between specific residue types. (B) Comparison of contact frequencies for residue type pairs. Black bars represent unoxidized (CTD) conditions, and red bars show oxidative conditions (CTD$_{ox}$). Error bars indicate the standard error of the mean.

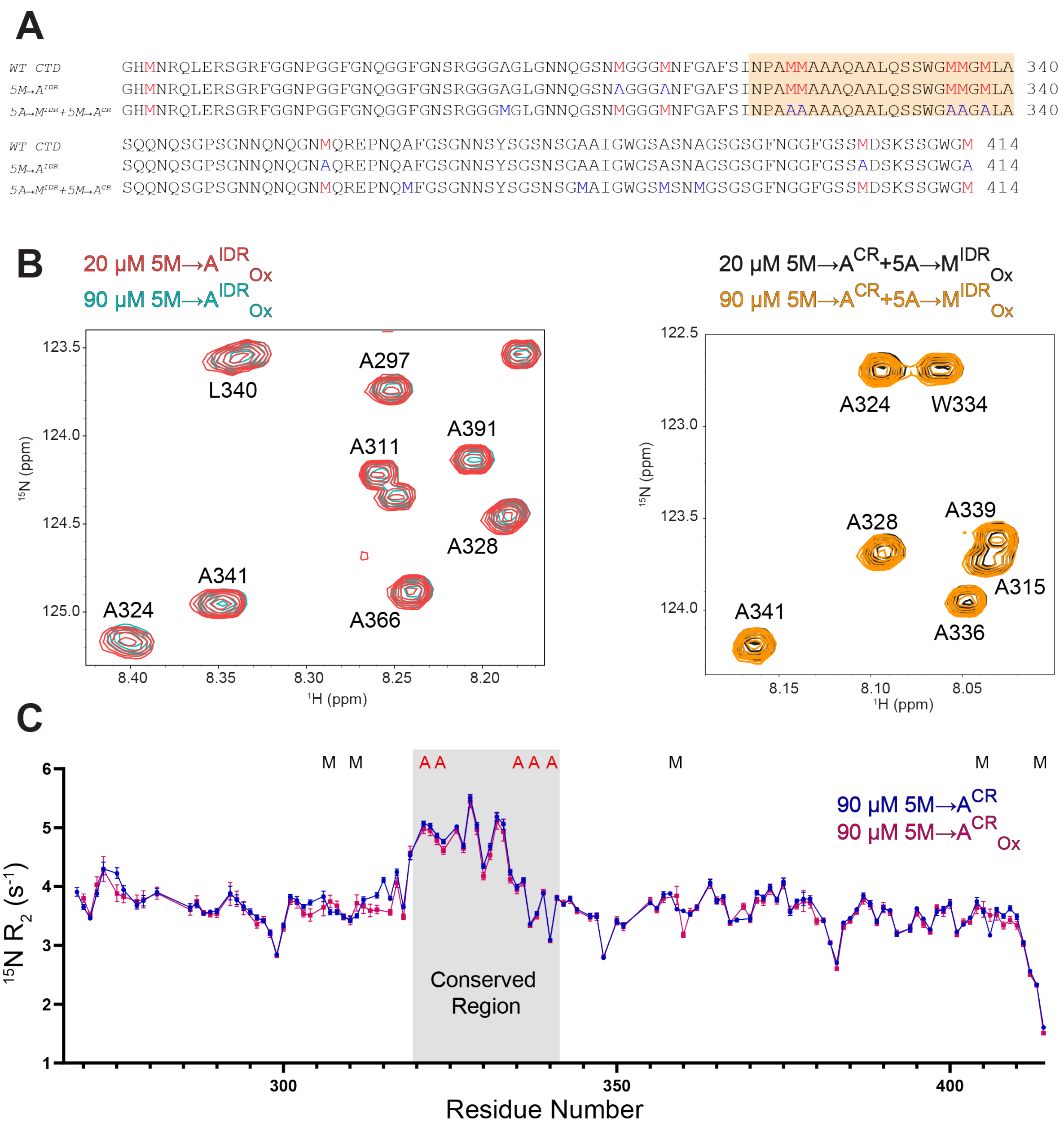

**Figure S8. Oxidation disrupts interactions of both methionines in the helical conserved region and within the disordered regions. (A)** Multiple sequence alignment of WT CTD as well as 5M→A$^{IDR}$ CTD and 5M→A$^{CR}$+5A→M$^{IDR}$ CTD used to probe the effect of oxidation on methionine interactions in structured versus unstructured regions. Position of natural methionines are denoted in red while positions of engineered alanines or methionines are denoted in blue. The structured conserved region is highlighted in orange. **(B)** Overlay of $^{1}$H-$^{15}$N HSQC spectra of 20 μM (light red) and 90 μM (cyan) oxidized 5M→A$^{IDR}$ CTD and 20 μM (black) and 90 μM (orange) oxidized 5M→A$^{CR}$+5A→M$^{IDR}$ CTD displaying that both constructs display no strong CSPs with increasing concentration following oxidation. **(C)** $^{15}$N $R_2$ analysis of unoxidized (dark blue) and oxidized (violet) 5M→A$^{CR}$ CTD demonstrating that oxidation of methionine residues in the disordered regions results in slight changes to the structural rigidity of the CTD near the methionines, including evidence for slightly faster motions surrounding M307 and M311, consistent with some contact disruption.

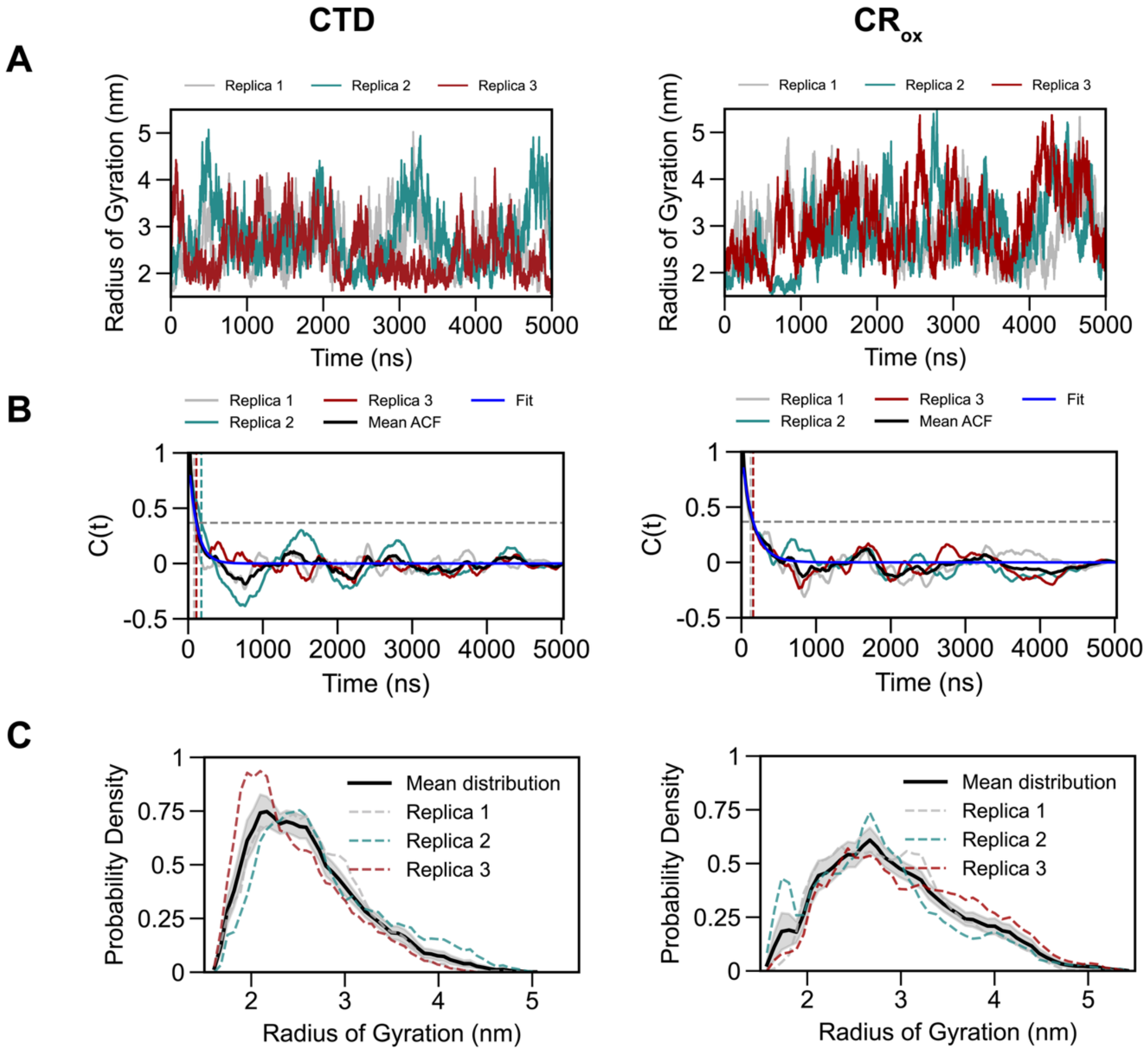

**Figure S9. Assessing the extent of conformational sampling across multi-replica trajectories of $IDR_{ox}$ and $CR_{ox}$. (A)** Rg variation as a function of time over three independent replica trajectories of $IDR_{ox}$ and $CR_{ox}$. **(B)** The plots show the time evolution of the radius of gyration autocorrelation function C(t) for each variant with data from three independent replicas. C(t) was calculated from the backbone radius of gyration (Rg) for each individual trajectory. The vertical dotted lines indicate where the autocorrelation function crosses 1/e (≈ 0.3679). Based on these crossing times, we conservatively selected 500 ns as the equilibration time for all systems. After this period, the autocorrelation functions consistently fluctuate around zero, indicating decorrelated sampling suitable for subsequent analysis. **(C)** Probability distributions of radius of gyration (Rg) with data from three independent replicas (shown in gray, blue, and red dashed lines) and their mean distribution (solid black line).

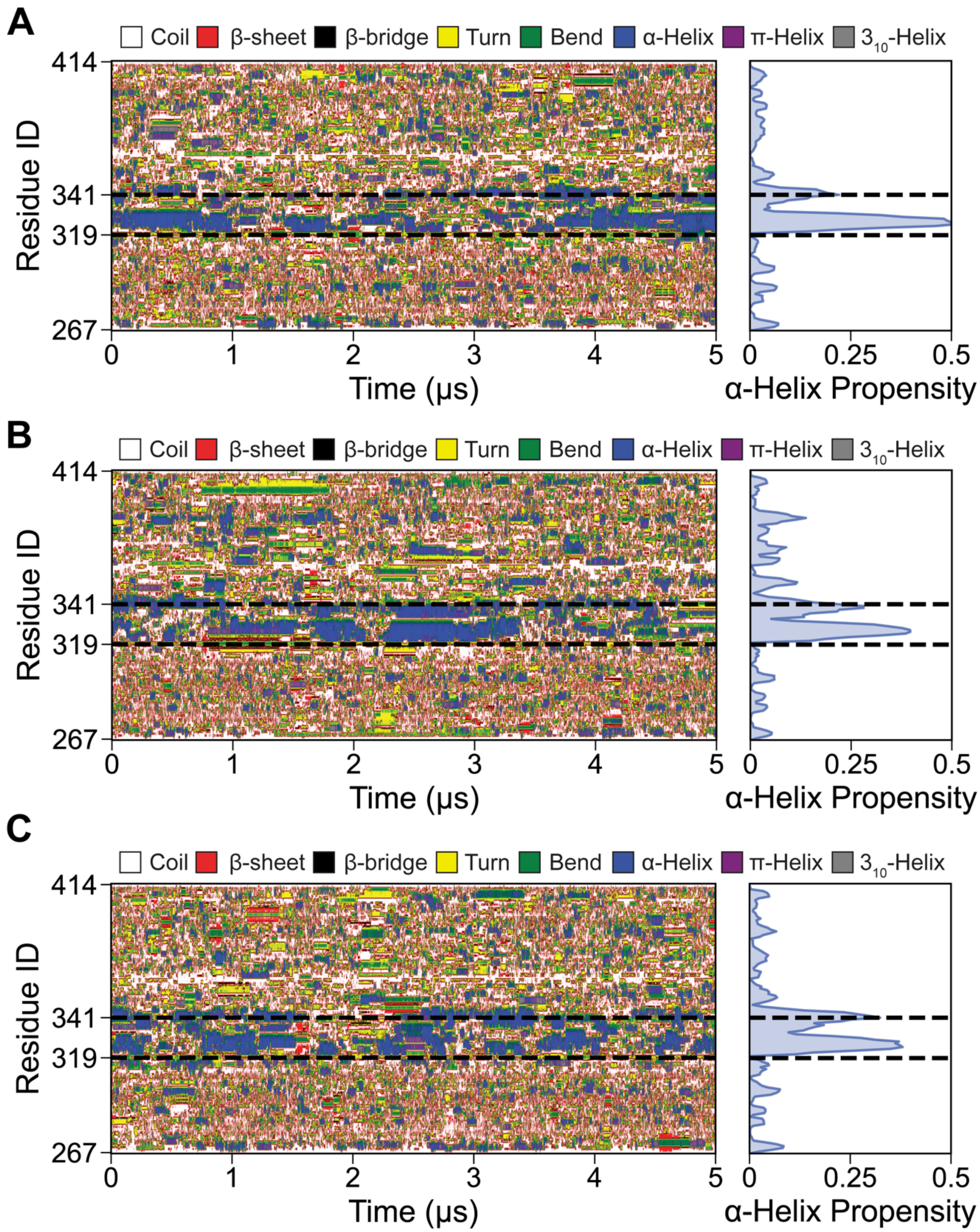

**Figure S10. Time evolution of secondary-structure changes upon flanking region Met oxidation. (A)** Replica 1, **(B)** Replica 2, and **(C)** Replica 3 showing residue-level secondary structure transitions over 5 μs.

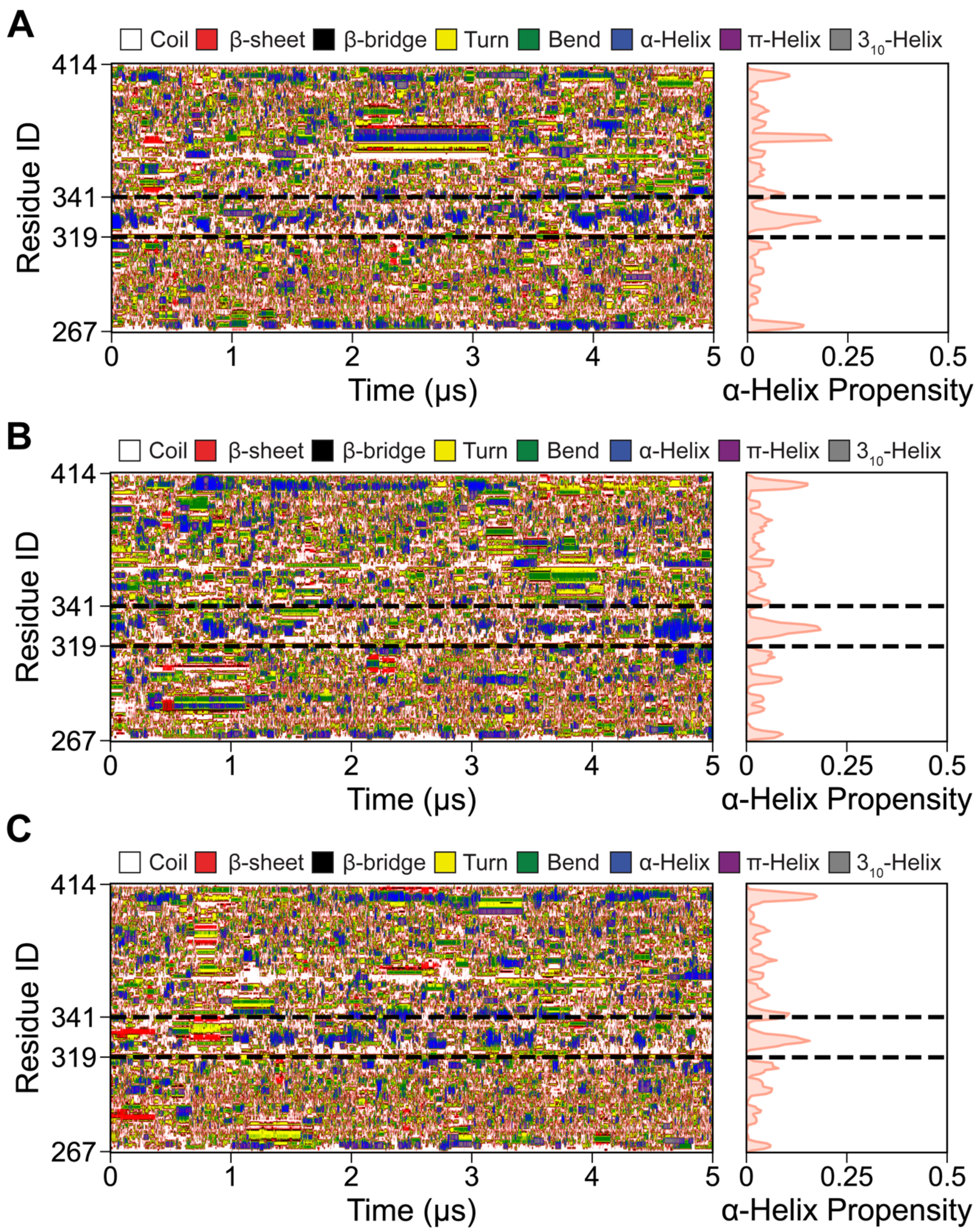

**Figure S11. Time evolution of secondary-structure changes upon CR Met oxidation**. **(A)** Replica 1, **(B)** Replica 2, and **(C)** Replica 3 showing residue-level secondary structure transitions over 5 μs.

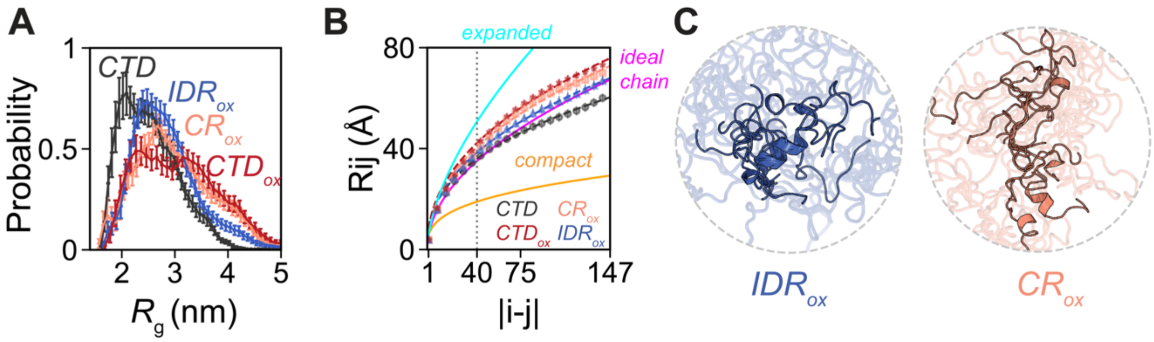

**Figure S12. Impact of IDR and CR oxidation on the CTD conformational ensemble. (A)** Probability distribution of the Radius of Gyration ($R_g$) for unoxidized (CTD, black), completely oxidized ($CTD_{ox}$, red) and position-dependent oxidized CTD variants ($IDR_{ox}$, blue; $CR_{ox}$, coral). **(B)** Root mean square intrachain distances for unoxidized and oxidized variants. **(C)** Representative structural ensembles of the $IDR_{ox}$ and $CR_{ox}$ variants.

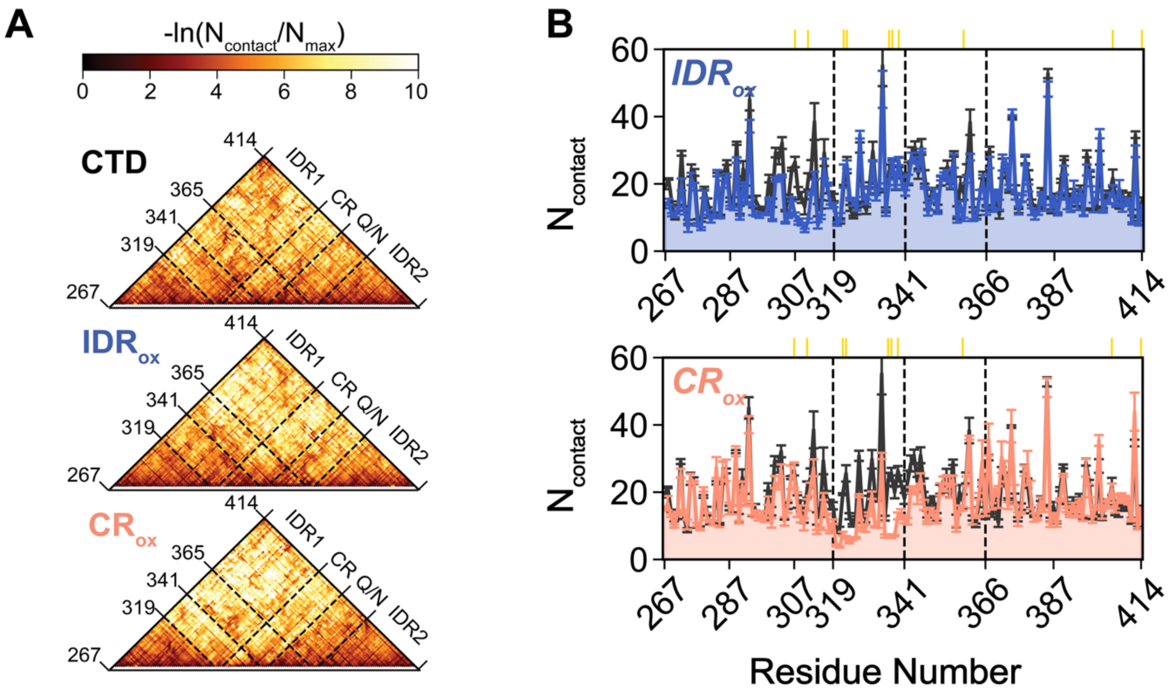

**Figure S13: Intramolecular Interaction Profiling upon Methionine Oxidation in Distinct Regions. (A)** Contact maps revealing intra-protein interactions in CTD (top), $IDR_{ox}$ (middle), and $CR_{ox}$ (bottom) conditions. Contact frequencies are represented as $-\ln(N_{contact}/N_{max}^{CTD})$, where darker red indicates higher contact probability. Region boundaries (IDR1, CR, Q/N, IDR2) are marked with dashed lines. **(B)** One-dimensional projection of the average residue-residue contact maps, quantifying the total contacts per residue for $IDR_{ox}$ (top), and $CR_{ox}$ (bottom). Error bars represent standard error of mean over three replicates for each construct.

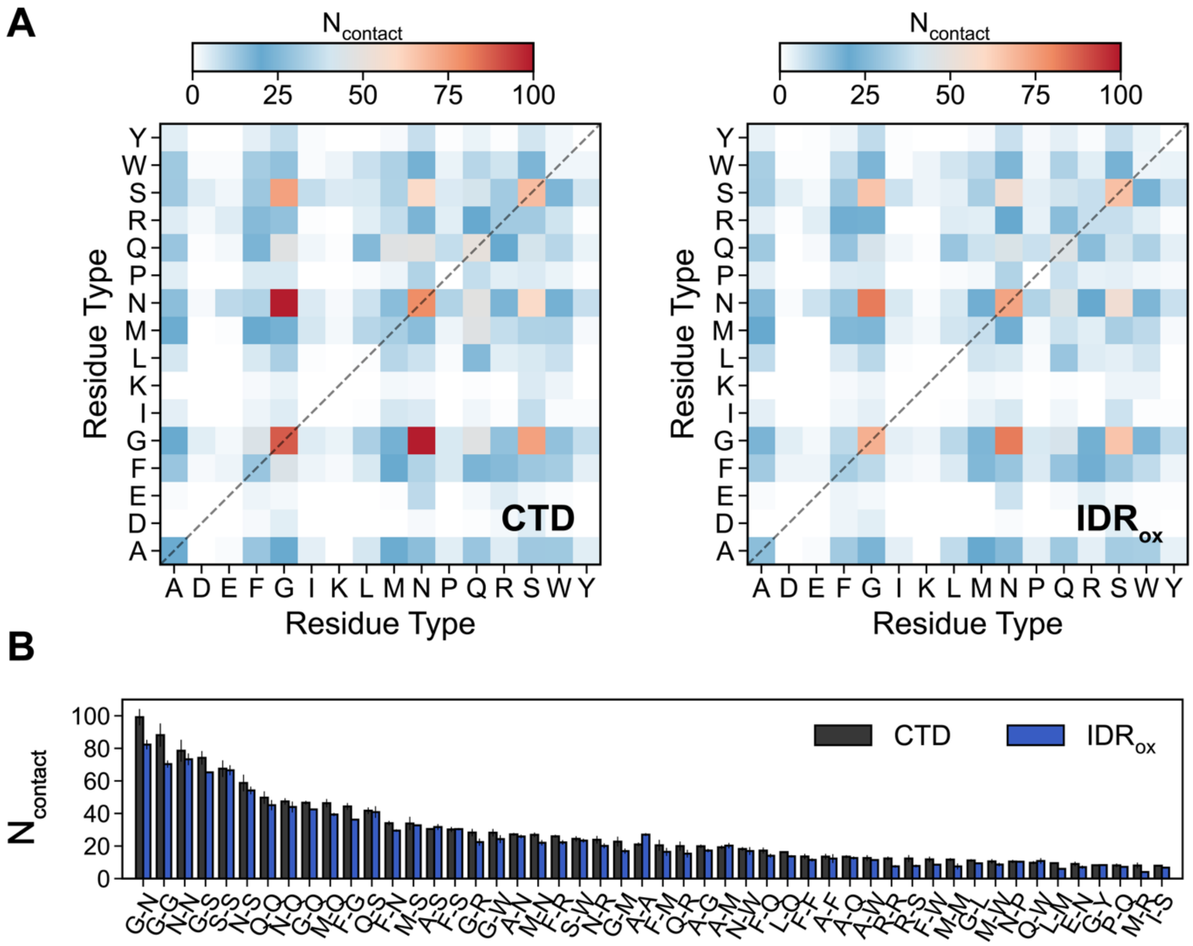

**Figure S14: Comprehensive Residue Type Interaction Analysis for IDR$_{ox}$ (A)** Interaction heatmap displaying contact frequencies between different amino acid types for CTD and upon oxidation of Met residues in flanking regions (IDR$_{ox}$). The color scale ranges from blue (low contact) to red (high contact), with the intensity indicating the number of interactions between specific residue types. **(B)** Comparison of contact frequencies for residue type pairs. Black bars represent unoxidized (CTD) conditions, and red bars show oxidative conditions (IDR$_{ox}$). Error bars indicate the standard error of the mean.

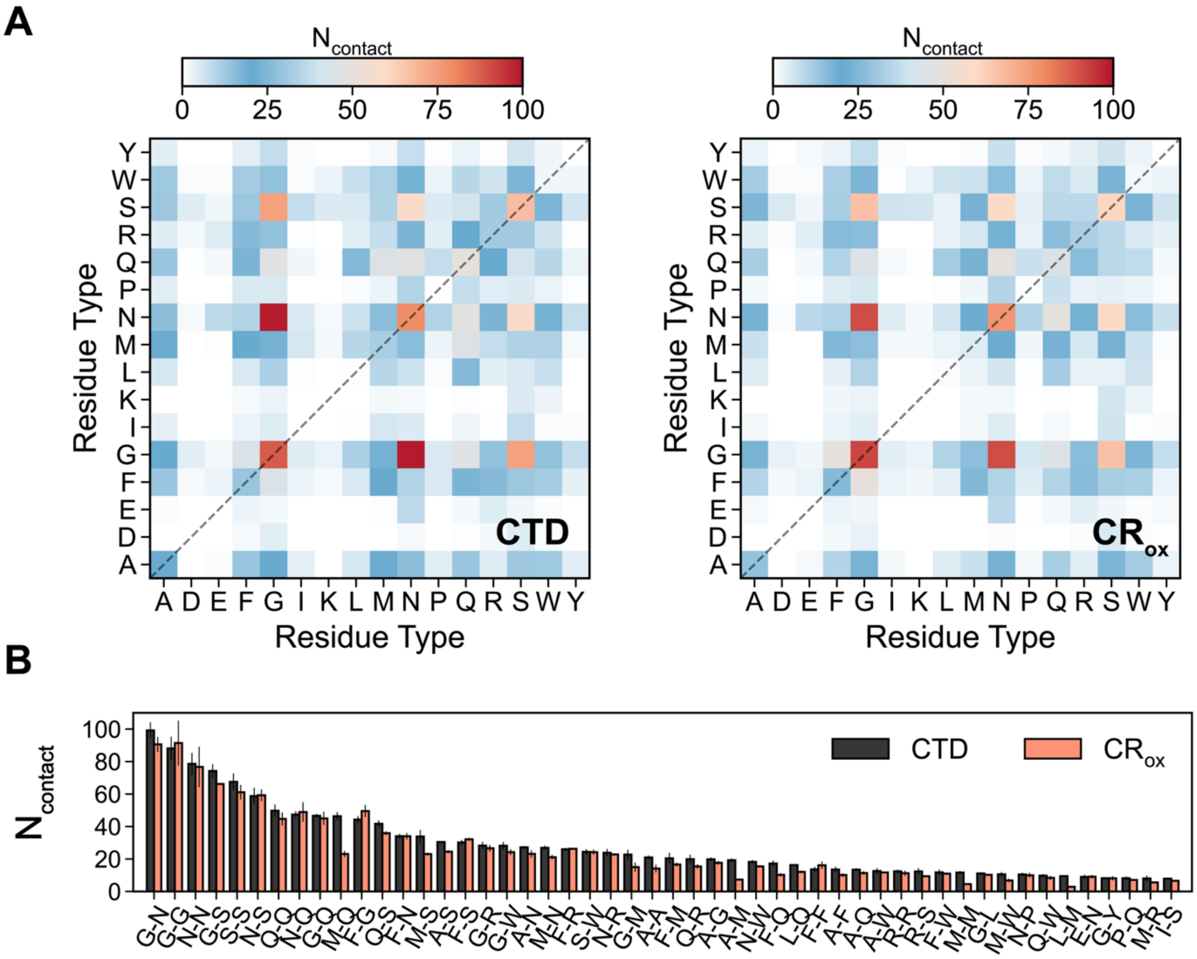

**Figure S15: Comprehensive Residue Type Interaction Analysis for CR$_{ox}$ (A)** Interaction heatmap displaying contact frequencies between different amino acid types for CTD and upon oxidation of Met residues in conserved region (CR$_{ox}$). The color scale ranges from blue (low contact) to red (high contact), with the intensity indicating the number of interactions between specific residue types. **(B)** Comparison of contact frequencies for residue type pairs. Black bars represent unoxidized (CTD) conditions, and red bars show oxidative conditions (CR$_{ox}$). Error bars indicate the standard error of the mean.

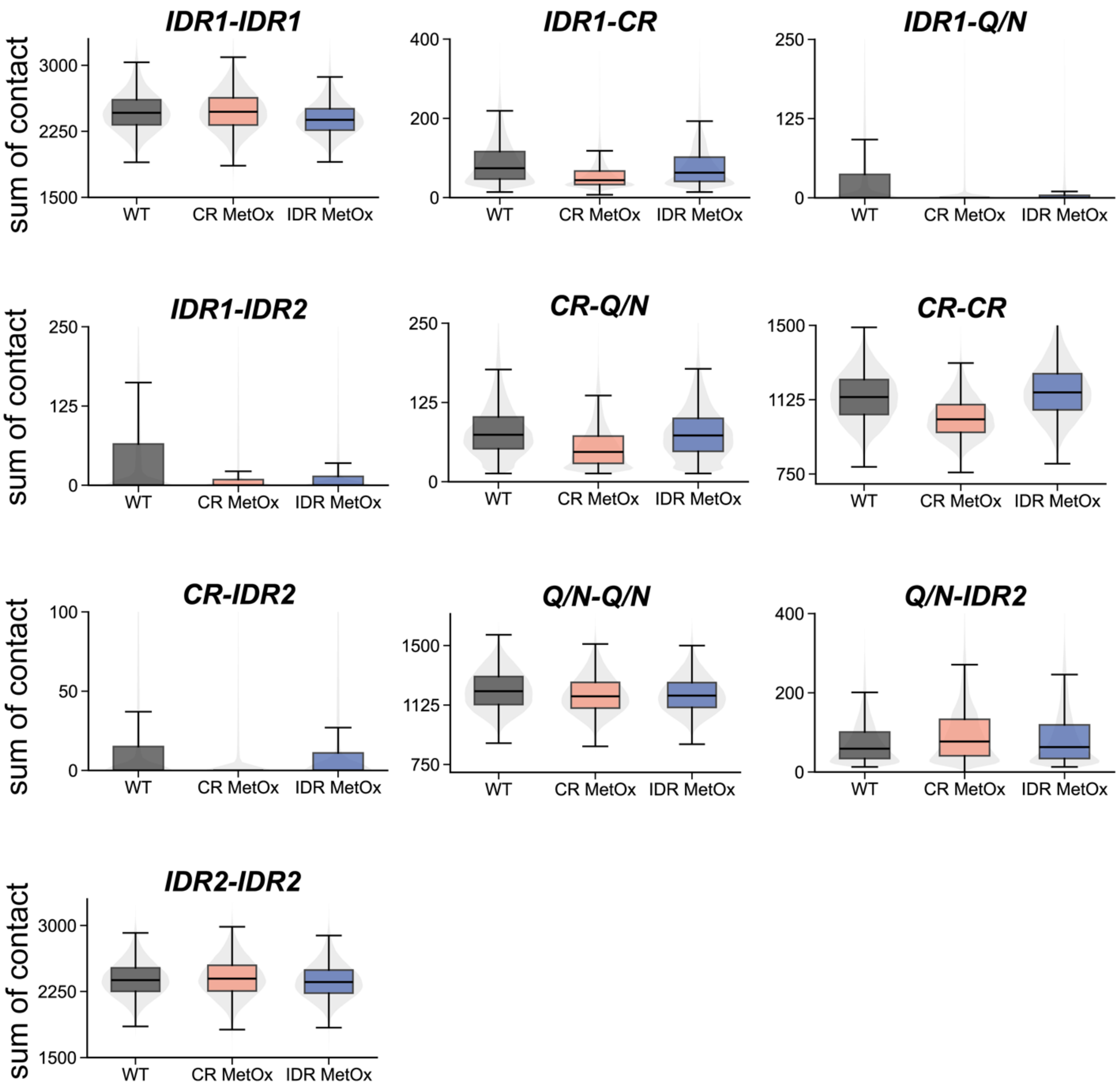

**Figure S16. Distribution of intra- and inter-region interaction contacts in CR$_{ox}$ and IDR$_{ox}$ constructs.** For each frame, all heavy atom contacts (4.5 Å cutoff) between protein residue pairs were calculated and grouped by region: IDR1 (residues 267-318), CR (residues 319-341), Q/N (residues 342-365), and IDR2 (residues 366-414). Both intra-region and inter-region interactions are shown. Box plots display medians (solid black lines) and means (dashed black lines), with boxes extending from first (Q1) to third (Q3) quartiles. Gray violin plots in the background reveal the full probability distribution of contact frequencies across all analyzed frames, highlighting differences in interaction patterns between CTD (gray), CR$_{ox}$ (orange), and IDR$_{ox}$ (blue).

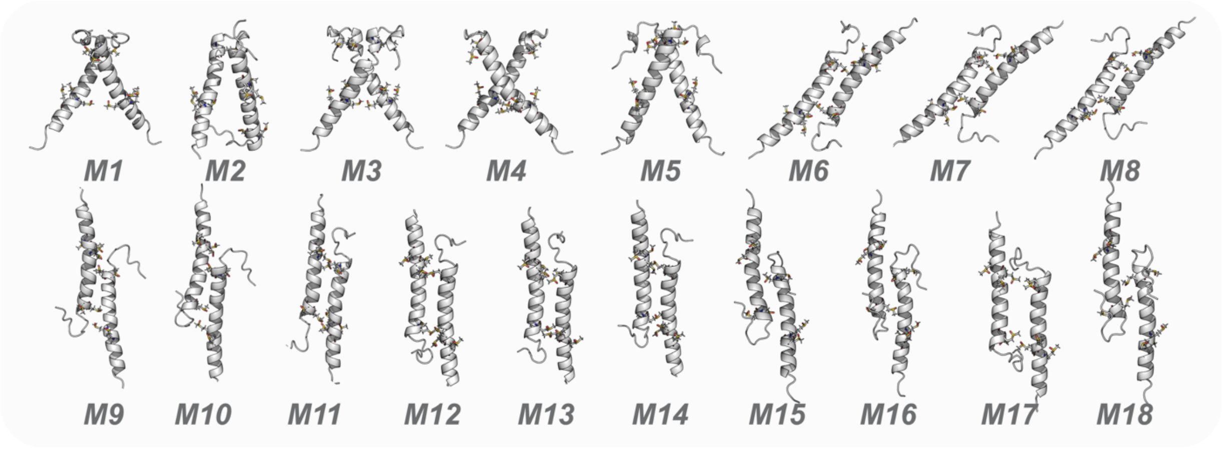

**Figure S17: Initial configurations for dimer simulations**. Models generated by Alphafold v2.3 in our previous study (Rizuan et al. (2025)[1]).

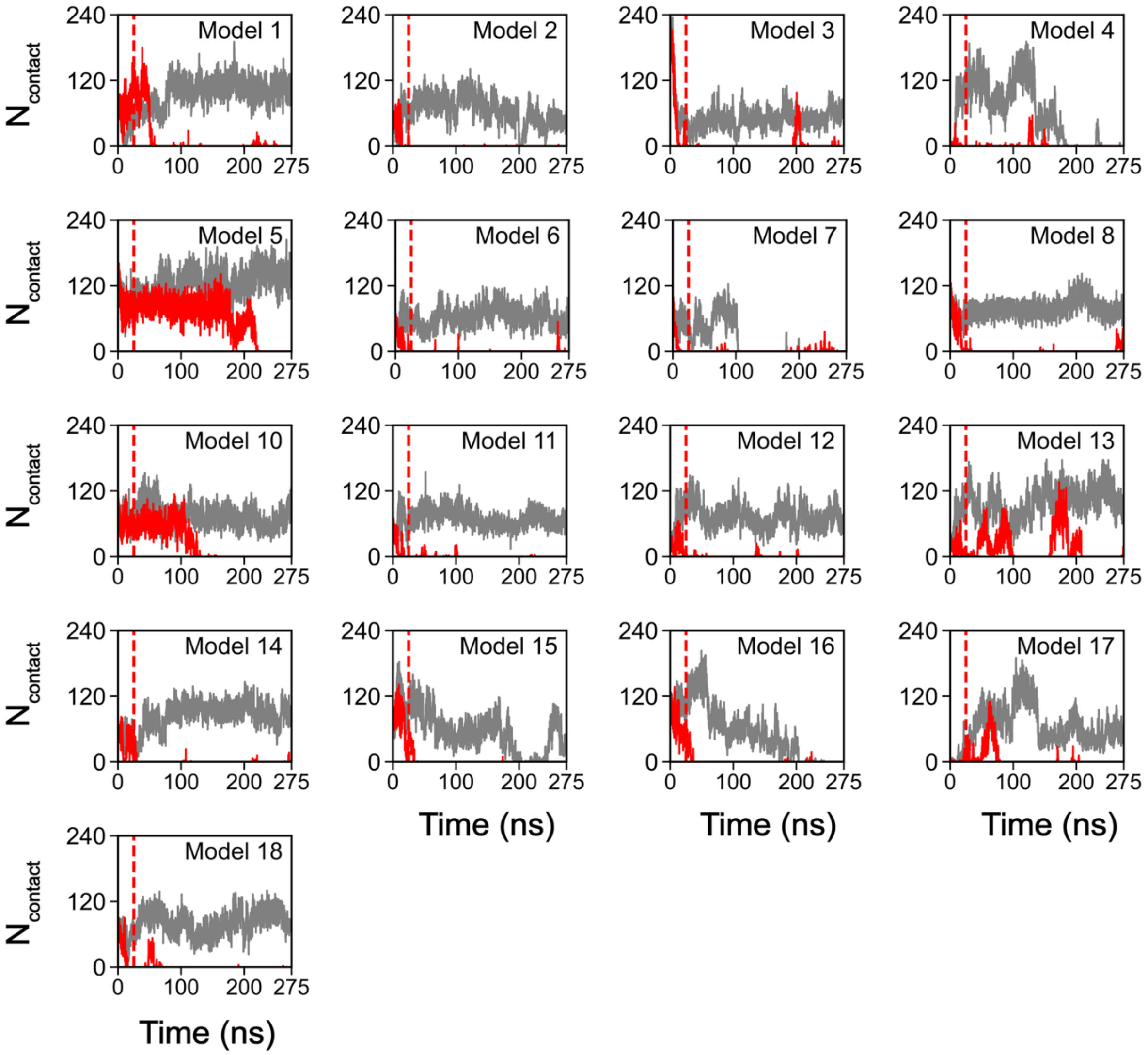

**Figure S18: Timewise Evolution of Contacts in Conserved Region (CR) Interface.** Gray solid lines represent unoxidized CTD and red solid lines represent dimers having Met residues oxidized in conserved region. Red dashed lines show the equilibration period (25 ns) prior to production runs (250 ns).
